# Eukfinder: a pipeline to retrieve microbial eukaryote genomes from metagenomic sequencing data

**DOI:** 10.1101/2023.12.28.573569

**Authors:** Dandan Zhao, Dayana E. Salas-Leiva, Shelby K. Williams, Katherine A. Dunn, Andrew J. Roger

**Author notes:** These authors contributed equally to this work (Dandan Zhao, Dayana E. Salas-Leiva).

## Abstract

Whole-genome shotgun (WGS) metagenomic sequencing of microbial communities allows us to discover the functions, physiologies, and evolutionary histories of microbial prokaryote and eukaryote members of diverse ecosystems. Despite their importance, metagenomic studies of microbial eukaryotes lag behind those of prokaryotes, due to the difficulty in identifying and assembling high-quality eukaryotic genomes from WGS data. To address this problem, we have developed Eukfinder, a bioinformatics pipeline that recovers and assembles nuclear and mitochondrial genomes of eukaryotic microbes from WGS metagenomics data. As part of its workflow, it utilizes two specialized databases to classify reads based on taxonomy which can be customized to the dataset or environment of interest. We applied Eukfinder to human gut microbiome WGS metagenomic sequencing data to recover genomes from the protistan parasite *Blastocystis* sp., a highly prevalent colonizer of the gastrointestinal tract of humans and animals. We tested Eukfinder using both a series of simulated gut microbiome datasets, which included varying numbers of *Blastocystis* reads combined with bacterial reads and by using real metagenomic gut samples containing *Blastocystis.* We compared the results of Eukfinder with other published workflows. With sufficient reads, Eukfinder efficiently assembles high-quality near-complete nuclear and mitochondrial genomes from diverse *Blastocystis* subtypes from metagenomic data without the aid of a reference genome. Furthermore, with sufficient depth of sequence sampling, Eukfinder outperforms similar tools used to recover eukaryotic genomes from metagenomic data. Eukfinder will be a useful tool for reference-independent and cultivation-free study of eukaryotic microbial genomes from environmental metagenomic sequencing samples.

**IMPORTANCE:** Rapid advancements in next-generation sequencing technologies have made whole-genome shotgun (WGS) metagenomic sequencing an efficient method for *de novo* reconstruction of microbial genomes from samples taken from different environments. So far, thousands of new prokaryotic genomes have been characterized from strains or species that were unknown to science. However, the relatively large size and complexity of protistan genomes has, until recently, precluded the use of the WGS metagenomic approach to sample microbial eukaryotic diversity. The bioinformatics pipeline we developed, Eukfinder, can recover eukaryotic microbial genomes from environmental WGS metagenomic samples. By retrieving high-quality protistan genomes from diverse metagenomic samples, we can increase numbers of reference genomes available to aid future metagenomic investigations into the functions, physiologies, and evolutionary histories of eukaryotic microbes in the gut microbiome and a variety of other ecosystems.

## INTRODUCTION

Microbial eukaryotes are ubiquitous and inhabit every global ecosystem. Their genomes can inform us of their physiological capacities, evolutionary histories, as well as their interactions with other microbes, their hosts, and their environment. Unfortunately, due to a lack of published complete genome sequences, high-throughput analyses of the population genomics of microbial eukaryotes have lagged behind those of prokaryotes. Prokaryotic genome data have dramatically increased in recent years thanks in part to the use of metagenomic assembly and binning tools, which allows for the study of uncultured microbial lineages (1, 2). Since the first application of a metagenomic sequencing approach to reconstruct near-complete bacterial genomes (3), thousands of high-quality complete or draft metagenome-assembled genomes (MAGs) for bacteria and archaeal species have been constructed (4, 5). However, due to the complex features of genomes from eukaryotic microbes (e.g., large size, increased sequence complexity, and a lower abundance in metagenomic sequencing data), the application of bioinformatic tools and workflows for generating eukaryotic MAGs is not well-established. This is mainly because typical metagenomics tools are either prokaryote-centered or too cumbersome to use, especially for researchers who lack a depth of bioinformatic knowledge (6).

To date, only a handful of investigations have used a metagenomic approach to reconstruct eukaryotic genomes. For example, Beghini and colleagues applied a bioinformatic method of reference-genome-based read mapping and assembly (hereafter referred to as ‘Refmapping’) to construct 43 draft *Blastocystis* genomes from 2154 gut metagenomic datasets. Amongst these genomes, 19 had sizes > 5 Mb, with completeness estimates ranging from 33% to 85%, based on the assembly size estimation (7). West and colleagues developed a machine learning k-mer-based tool, EukRep, for separating eukaryotic and prokaryotic sequences (8). A recent study applied EukRep to 1198 metagenomic datasets and in total 14 novel eukaryotic genomes were recovered with estimated median completeness of 91% (9). Tiara, which uses a deep learning approach (a subfamily of machine-learning techniques), was introduced to identify eukaryotic and organellar genomes within metagenomic datasets (10). These studies demonstrate that it is possible to reconstruct microbial eukaryotic genomes from metagenomics datasets.

Each of the previous pipelines however have limitations. For example, the reference genome mapping method will not be useful for recovering genomes in metagenomic data unless there are closely related reference genomes available. Additionally, the performance of the machine-learning approaches EukRep and Tiara, can be affected by the training reference genome sets utilized, and adjusting these models to construct custom databases would require retraining of the models. Furthermore, these models are designed ideally to work on contigs >3000 bp in length.

To circumvent these limitations, we developed Eukfinder, a bioinformatic tool for recovery and assembly of eukaryotic nuclear and mitochondrial genomes from environmental metagenomes. Eukfinder improves upon existing pipelines by allowing for the use of short reads and the ability to customize the search database. Specialized databases can be built by users to include reference genomes from representative organisms in the environment of interest. To demonstrate its utility, Eukfinder was applied to mock community datasets that included varying amounts of *Blastocystis* sequence reads as well as to real human gut metagenomic datasets. We examined the ability of Eukfinder to identify eukaryotic sequences and assemble nuclear and mitochondrial genomes from those sequences, focusing on *Blastocystis* genomes as a test case.

*Blastocystis* spp. (referred to as *Blastocystis* hereafter) are amongst the most prevalent microbial eukaryotes colonizing the GI tracts of different animals and humans (11). Out of 17 currently known subtypes (STs), only 8 have published genomes (Table S1) (12). Among these genomes, three (ST1, ST4, and ST7) are nearly complete (13–15) whereas the remainder are rough draft assemblies (16). Here, we show that applying Eukfinder to metagenomic data facilitates the reconstruction of genomes of microbial eukaryotes like *Blastocystis*. Eukfinder recovers more complete nuclear and mitochondrial genomes than both reference mapping and machine learning-based methods. Our new approach to identifying and assembling eukaryotic genomes from metagenomic data will help in increasing the number of reference genomes available, and pave the way for future studies into the function, physiology, and evolutionary history of microbial eukaryotes.

## RESULTS

### Overview of workflow

Eukfinder is designed to retrieve eukaryotic genomes, both nuclear and mitochondrial, from metagenomic data (Fig 1). Eukfinder can use either short Illumina metagenomic reads (processed with Eukfinder_short), assembled contiguous sequences (hereafter referred to as contigs), or metagenomic reads derived from long-read sequencing platforms, such as Oxford Nanopore or PacBio (processed with Eukfinder_long). Eukfinder_short processes short reads first by classifying them into 5 distinct taxonomic categories (Archaeal, Bacterial, Viral, Eukaryotic, and Unknown) using Centrifuge (17), a read classification engine that uses a custom database (DB1), and PLAST (18), a more sensitive sequence similarity search tool, with a second specialized database (DB2) (Fig 1a). Here, because we were focused on retrieving eukaryotic microbes from gut metagenomic data, DB1 was built to maximize the representation of microbes from the gut (full details on the assembly of databases are given in the **MATERIALS AND METHODS**). DB2, while having similar taxonomic representation as DB1, was smaller to ease computational burden. Following the first round of classification, all reads identified as ‘Eukaryotic’ or ‘Unknown’ were automatically assembled into contigs using metaSpades by Eukfinder.

**Figure 1.**
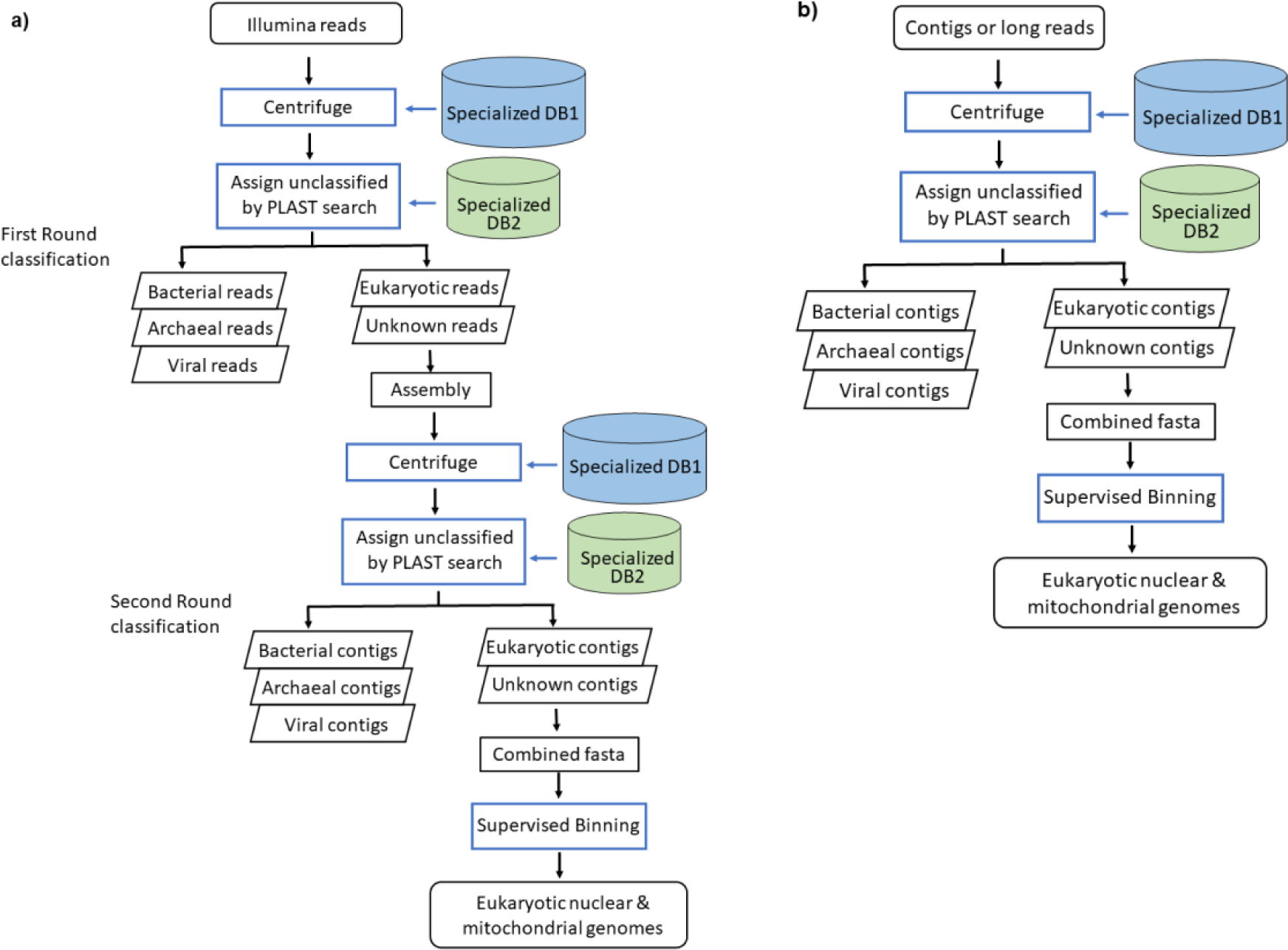
Schematic representation of Eukfinder workflows. Eukfinder is a taxonomic classification-based bioinformatics approach to retrieve microbial eukaryotic nuclear and mitochondrial genomes from WGS metagenomic sequencing data. Eukfinder has two different workflows based on the input files, a) Eukfinder_short using Illumina short reads, it first classifies Illumina reads into 5 distinct taxonomic categories (Archaeal, Bacterial, Viral, Eukaryotic, and Unknown), after assembling the Eukaryotic and Unknown reads together, second round of classification and supervised binning, it will output Eukaryotic nuclear and mitochondrial genomes, and b) Eukfinder_long using assembled contigs or long-read sequencing data generated by Nanopore or Pacbio platforms. It goes through one round of classification to select Eukaryotic and Unknown contigs, and after supervised binning generates Eukaryotic nuclear and mitochondrial genomes.

Assembled contigs were then classified with Eukfinder using Centrifuge and PLAST with their associated databases. Contigs assigned as ‘Eukaryotic’ or ‘Unknown’ were subjected to supervised binning using depth of coverage and taxonomy for bin refinement. The final outputs were high quality draft assemblies of nuclear and mitochondrial eukaryotic sequences. If starting with contigs or long-read sequences as input data, then the Eukfinder_long workflow (Fig. 1b) is used. The latter is similar to Eukfinder_short however it only has one round of classification through Centrifuge and PLAST. Full details of the workflow and parameters are given in the **MATERIALS AND METHODS**.

To test Eukfinder’s ability to retrieve *Blastocystis* genomes and examine its performance relative to other eukaryote metagenomic read classifiers, we benchmarked Eukfinder_short and Eukfinder_long along with three other methods on simulated mock community datasets. These controlled metagenomic datasets consisted of synthetic human gut bacterial data (16.6M reads) and varying amounts of randomly-selected *Blastocystis* ST1 Illumina sequencing reads (from 300K to 10M reads, hereafter referred to as ‘Randomly-selected’ *Blastocystis* reads), which resulted in eight groups of mock metagenomic datasets in quadruplicate (Table S2) (see **MATERIALS AND METHODS** for more details).

### Identifying optimal parameters for alternative methods

Due to differences in input data for the methods examined, Eukfinder_short, and Refmapping (7) were used to examine short Illumina reads, while Eukfinder_long, EukRep (8) and Tiara (10) were used to examine assembled contigs of these reads. To ensure the comparisons were fair, we first identified the parameters for each program that maximized genome completeness and contiguity. The following section summarizes our efforts to find the best parameters for Refmapping, EukRep, and Tiara.

Refmapping can use two different alignment strategies to map reads, global (end-to-end) or local mapping. We examined the performance of both under three test groups of mock community datasets (500K, 1M, 2M of random selected *Blastocystis* reads) and found that local alignments generated more complete and contiguous genomes compared to global (end-to-end) alignments (Fig. S1a). This was especially true when there were lower numbers of reads to map to the reference genome. With 500 K reads, the genomes recovered using the local alignment had an average completion of >60% while the genomes recovered using global alignment were <20% complete. When the number of *Blastocystis* reads increased to 1M, the genomes recovered using the local alignment had an average completion of 90% while the genomes recovered using global alignment were <60% complete. Therefore, we ran Refmapping with local alignment on all datasets and used the *Blastocystis* ST1 genome as the reference sequence for the mock community and ST3 and ST4 genomes for the human metagenome data, for comparison with Eukfinder_short.

EukRep allows for differences in algorithm stringencies when identifying matches. We examined all three algorithm stringencies (strict, balanced, and lenient) and found that the lenient stringency provided the most complete genome recovery at all read counts tested (Fig. S1b). The strict mode, which was designed to have a lower false positive rate generated the least complete genomes and lenient modes the most complete genomes with balanced mode intermediate. On average, the balance mode increased the completeness by 0.5-1.5% more than the strict mode; the lenient mode further increased the completion 2-5%, while setting “tie” decision handling to prokaryotic, generated genomes 1% less complete than when the default (eukaryotic) was used. Based on these findings, we ran EukRep with lenient mode and default eukaryotic “tie” decision for all datasets for comparison with Eukfinder_long.

Tiara has two main parameters which can be adjusted in its workflow, the k-mer size, and probability threshold. To examine these parameters Tiara was tested under a combination of three different k-mers (4-mer, 5-mer, and 6-mer) and five different probability thresholds (p=0.5, 0.55, 0.6, 0.65, 0.7). We found that utilizing a k-mer size of 4 and probability threshold of 0.7 produced the most complete and contiguous genomes (Fig. S1c). Therefore, Tiara was run with a k-mer of 4 and a probability threshold of 0.7 for all datasets for comparison with Eukfinder_long.

### Mock community dataset analyses

To explore the ability of Eukfinder to recover eukaryotic genomes, we compared the short and long-read workflows with Refmapping, EukRep, and Tiara to recover *Blastocystis* ST1 genomes from the eight groups of mock metagenomic datasets (Fig. 2, Table S3). Completeness of the *Blastocystis* ST1 nuclear genome was evaluated based on total length of all recovered ST1 contigs (Fig. 2a), genome fraction (% of genome completion), N50 and L50 using QUAST for all methods (Fig S2). When comparing Eukfinder_short and Refmapping we found that Refmapping with optimal parameters (local mode and the ST1 reference genome) recovered the most complete genomes when datasets had low number of *Blastocystis* reads (≤1M) (Fig 2a, Fig. S2) The difference in size and completeness between Eukfinder_short and Refmapping was largest at lower read counts, with Refmapping recovering between ∼18% to 10% more complete genomes than Eukfinder_short at 300K to 500K reads (Fig. S3a-c). This gap decreases as the number of *Blastocystis* reads increases, with Refmapping recovering ∼3.5% to 1% more complete genomes at 750K and 1M *Blastocystis* reads respectively (Fig. S3d-e). When the number of *Blastocystis* reads was >1M, Eukfinder_short recovered statistically longer and more complete *Blastocystis* genomes than Refmapping (Fig 2a, Fig S2, Table S4, p<0.005).

**Figure 2.**
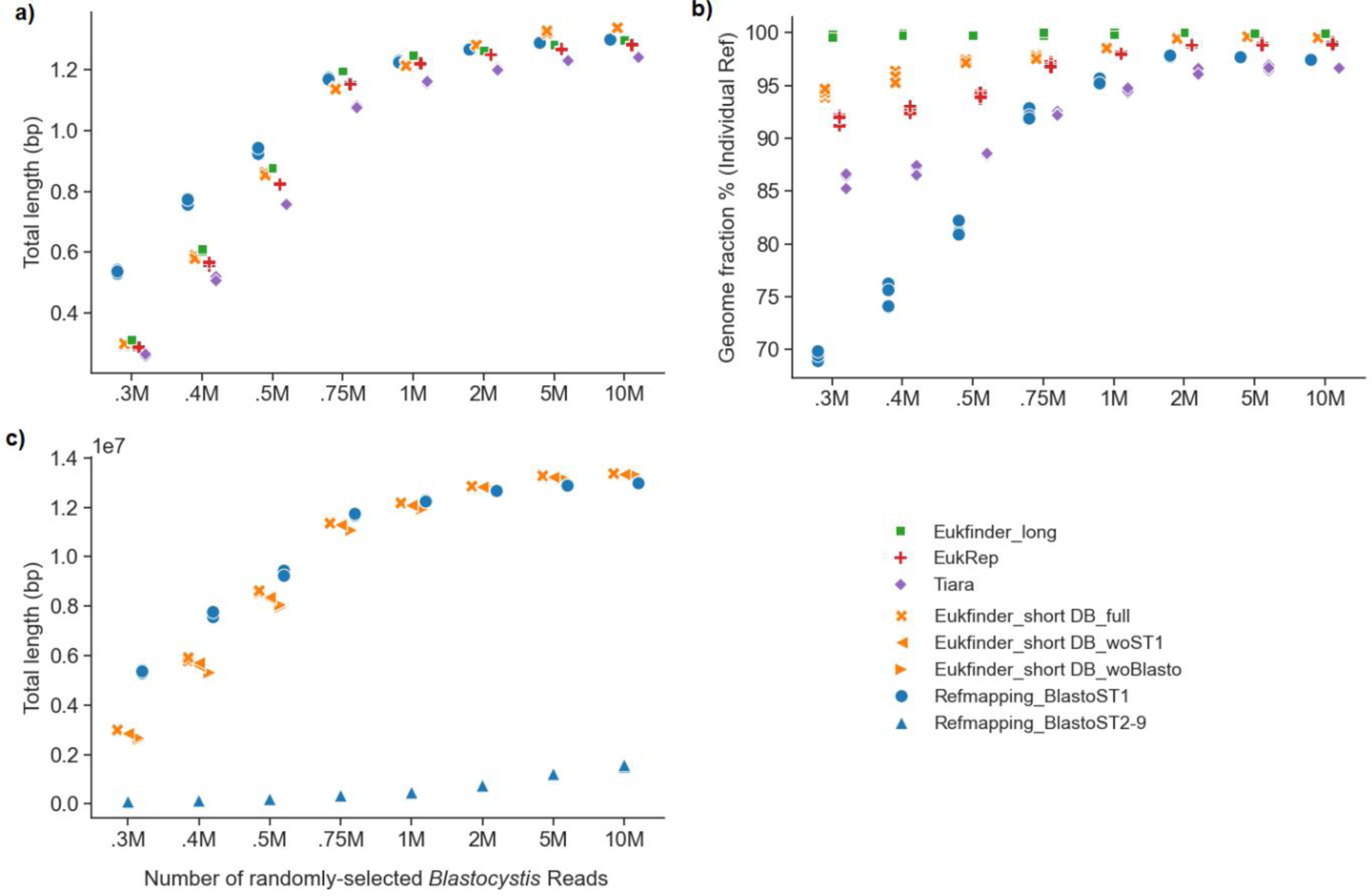
Comparison of methods (Eukfinder_short, Refmapping, Eukfinder_long, EukRep, and Tiara) to recover *Blastocystis* ST1 genomes in eight datasets consisting of bacterial mock metagenomic data (mix-51) with varying amounts of randomly selected *Blastocystis* ST1 Illumina sequencing reads. a) Plot of total length of *Blastocystis* ST1 contigs recovered by each method. b) The fraction of true *Blastocystis* contigs (Individual Ref) that were retrieved from each mock community dataset assessed using QUAST. Fraction of the randomly selected *Blastocystis* reads (IndRef) identified, assessed using QUAST. c) Plot of *Blastocystis* ST1 total length recovered using alternative reference datasets with Eukfinder_short and Refmapping. Eukfinder_short was assessed excluding the ST1 genome (woST1), and with no *Blastocystis* genomes (woBlasto) in the database. Refmapping was assessed using ST2-9 genomes as the reference (see supplemental table 1).

We used Eukfinder_long, EukRep and Tiara to compare total length and genome completion of the *Blastocystis* genome from classification of assembled contigs (≥1000 bp). *Blastocystis* genomes were significantly longer and more complete with Eukfinder_long than with EukRep (0.8-3.4%) or Tiara (2.9-8%) identified reads (Bonferroni corrected p<0.05 pairwise t-test, Fig. S2a, Table S5 & S6). Of the three methods using contig data, Tiara had shorter and less complete genomes than either EukRep or Eukfinder_long (Fig. 2a and 2b, Fig S2).

Precision and recall of the genomes recovered by the three methods was also examined. Precision examines how many of the contigs that a method identified as *Blastocystis* were in fact *Blastocystis* (true positives divided by true positives plus false positives) while recall is how well a method performed at recovering all of the *Blastocystis* contigs (true positives divided by true positives plus false negatives). The true *Blastocystis* contigs that could be retrieved from each mock community dataset were used as reference to evaluate the genomes recovered by each method using QUAST (Fig. 2b). Both Eukfinder_long and EukRep had high precision (∼100%, Figure 3), while precision for Tiara ranged from 96-99% depending on the number of *Blastocystis* reads. The high precision suggests that contigs identified as *Blastocystis* by the methods were correct. Recall on the other hand, which is the method’s ability to identify all of the *Blastocystis* contigs, varied much more. Eukfinder_long had nearly perfect recall in all cases (>99.7%) while EukRep recall ranged from 91-97%, increasing as the number of reads increased (Figure 3a). Tiara had the worst recall among the methods (85-92% Figure 3a). The lower rate of recall in EukRep and Tiara suggests that these methods are failing to identify all the *Blastocystis* contigs, in some cases missing up to 15% of the contigs (Figure 3b).

**Figure 3.**
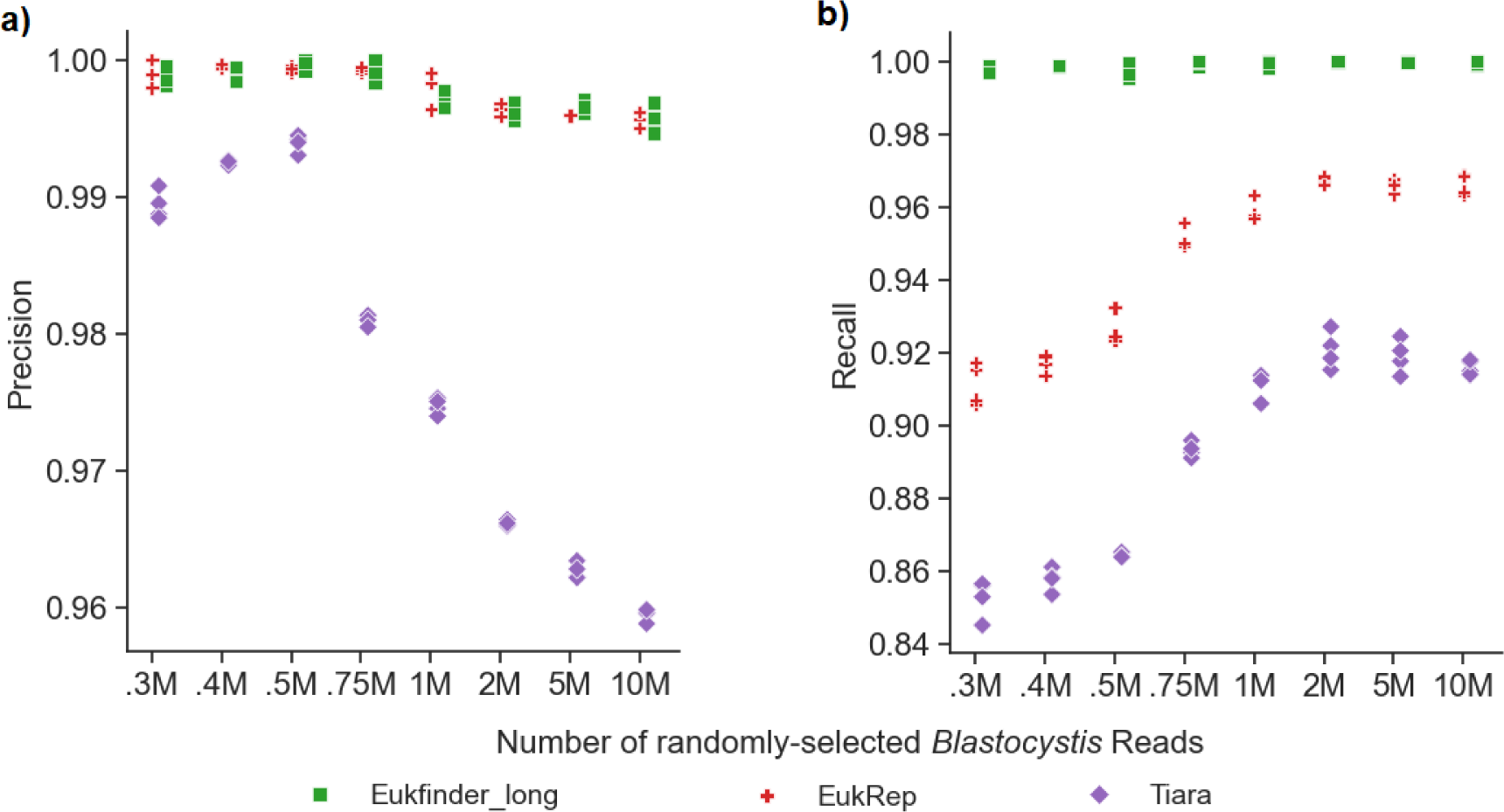
Precision and recall of the genomes recovered by Eukfinder_long, EukRep, and Tiara. a) Average precision, measured as the fraction of true *Blastocystis* reads among all reads assigned as *Blastocystis*, and b) average recall, measured as the fraction of *Blastocystis* reads that were recovered, of the four replicates at each read depth for the three long-read methods (Eukfinder_long, EukRep, and Tiara). Precision = true positives / (true positives + false positives). Recall = true positives / (true positives + false negatives).

### When there is no reference genome

Reference mapping methods do well when there is a good reference genome available, making this method dependent on the quality, and availability of a draft reference genome of the organism of interest. Likewise, Eukfinder also benefits if the reference genome is represented in its Centrifuge and PLAST databases. To examine the impact of the reference genome on our ability to recover genomes we replaced the *Blastocystis* ST1 genome with *Blastocystis* ST2-ST9 (ST2-9) as the reference for Refmapping and investigated two modifications to the databases for Eukfinder, one in which ST1 was not included (DB_woST1) but other *Blastocystis* genomes remained and one in which no *Blastocystis* genomes were included (DB_woBlasto). We found that Refmapping using ST2-9 as the references performed very poorly recovering < 12% of the *Blastocystis* ST1 genome and recovering shorter genomes regardless of number of reads included (Fig. 2c; Fig S3a), highlighting the impact of the reference genome on the ability of Refmapping to recover the genome. Eukfinder_short by contrast recovered genomes of similar total length in all cases with a <2% reduction in genome fraction recovered using the woST1 database and <4.3% reduction using the woBlasto database compared to when the specialized DBs were used (Fig 2c; Fig S3a). The reduction in genome completion when the *Blastocystis* ST1 genome was not included was much smaller (e.g., genome fraction <1 to <4.5%) with Eukfinder_short in comparison to the reduction in completion yielded by Refmapping (e.g., genome fraction 36 to 88%) (Fig 2c; Fig S3a).

### Real human gut metagenomic samples

Eight human gut metagenomic samples containing *Blastocystis* reads (Table S7) were examined using Eukfinder (short and long), Refmapping, EukRep and Tiara to assess the ability to recover *Blastocystis* reads. Four of the samples examined contained *Blastocystis* ST3 sequences (samples 3A-3D) and four samples contained ST4 sequences (samples 4A-4D). We assessed total genome length, genome fraction, N50 and %GC for the genome recovered from all samples using QUAST and examined the number of single copy genes (SCG) for each using BUSCO (20) with the Stramenophile dataset (21; stramenophile_odb10) (Fig 4; Fig S4 & S5; Table S8). Examining the performance of short read methods across all eight samples using pairwise comparisons we found that Eukfinder_short recovered in significantly longer genome lengths than Refmapping (p=0.013), with one exception (sample 4A, Fig 4). While Eukfinder_short had significantly longer total length than Refmapping, the genome fraction recovered, and the number of detected SCGs across samples did not differ significantly between them (p=0.20, and 0.06 respectively, Fig 4; Fig S4; Table S8). Examining long read methods Eukfinder_long recovered more complete genomes based on total length (vs EukRep p=7.6e-5; vs Tiara p=6.2e-5; Fig 4a & c; Table S8), fraction of genome recovered (vs EukRep p=1.6e-4; vs Tiara p=5.2e-5; Fig S4; Table S8) and number of SCG (vs EukRep p=2.0e-4; vs Tiara p=7.1e-4; Fig 4b & d; Table S8). *Blastocystis* genomes recovered by Eukfinder_long were 2 to 15% more complete than EukRep genomes and 6-20% more complete than Tiara genomes (Table S8).

**Figure 4.**
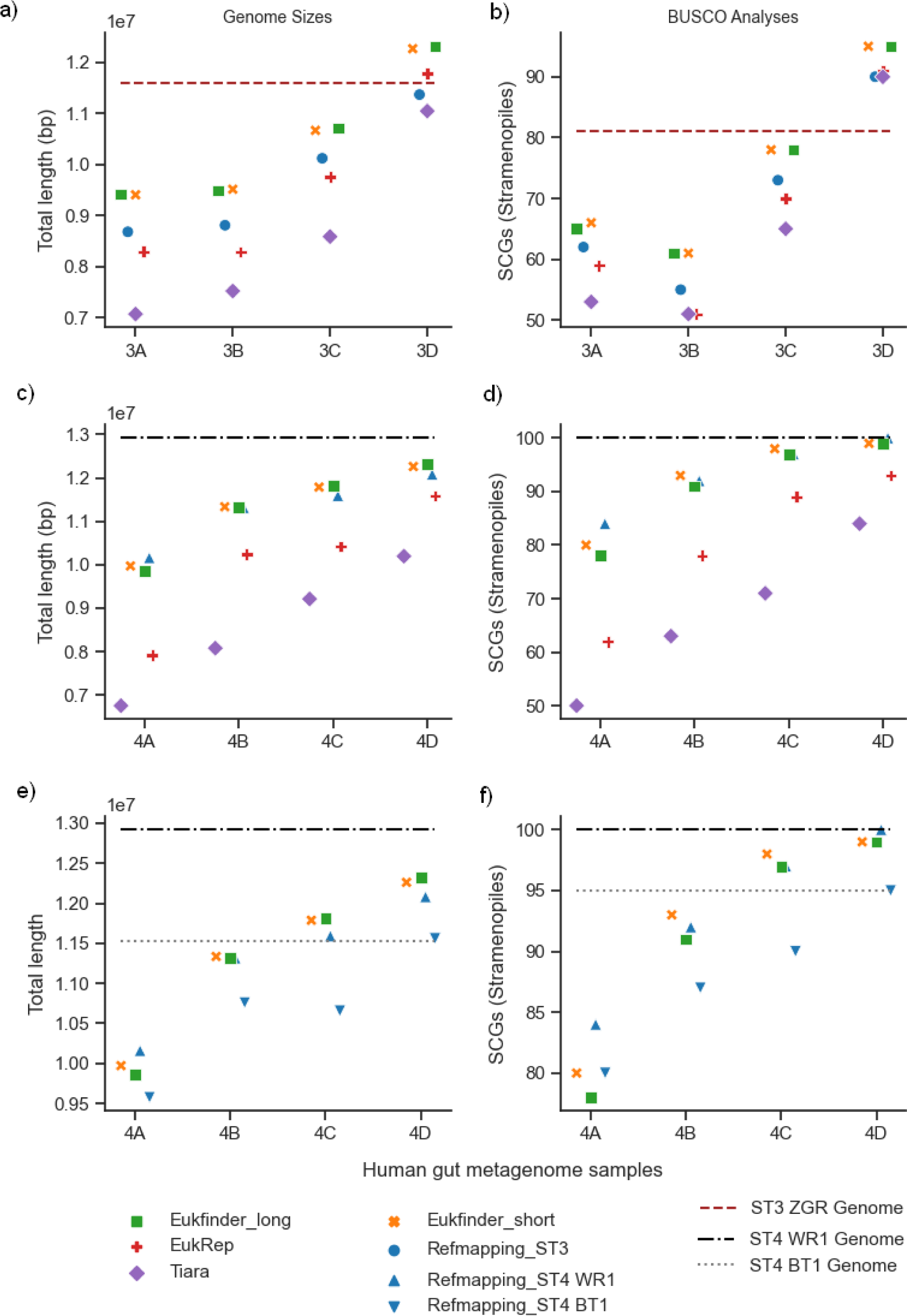
*Blastocystis* ST3 and ST4 genomes recovered from eight human gut metagenome samples (3A-D and 4A-D) using Refmapping, Eukfinder_short, Eukfinder_long, EukRep and Tiara. Genome completeness in ST3 containing samples (Refmapping reference genome ST3 ZGR) based on a) total length (bp) and b) number of single-copy genes (SCGs) detected using BUSCO (stramenophile_odb10). Genome completeness in ST4 containing samples (Refmapping reference genome ST4 WR1) based on c) total length (bp) and d) number of single-copy genes (SCGs) detected using BUSCO (stramenophile_odb10). Reanalysis of Refmapping ST4 genome completeness using reference genome ST4 BT1, e) total length (bp) and f) number of single-copy genes (SCGs) detected using BUSCO (stramenophile_odb10). Tiara was runwith kmer of 4 and probability threshold of 0.7, and Refmapping was run in local mode. The brown, and black dashed lines, and grey dot line represent the assessment results for *Blastocystis* ST3 ZGR, ST4 WR1 and ST4 BT1 reference genomes, respectively.

Among the samples containing *Blastocystis* ST3, short read methods recovered genomes that ranged in size from 8.7 to 12.3 Mbp and had 74 to 97% completion while long read methods recovered genomes between 7.5 Mb to 12.3 Mbp and had completion between 59 to 97% (Fig 4a-b; Fig. S4). One sample (3D) had a genome recovered by Eukfinder_long, Eukfinder_short and EukRep (12.315 Mb, 12.270 Mb, and 11.783 Mb respectively) that was longer than the available published reference genome (11.589 Mb) (Fig 4a; Table S1). In addition, all methods recovered more SCGs using BUSCO (stramenophiles_odb10) in sample 3D than the published reference genome (Fig 4b). This is likely because the ST3 reference genome was also recovered using a metagenomic method and was estimated as being incomplete.

Among the samples containing *Blastocystis* ST4, short read methods recovered genomes that ranged in size from 10 to 12.3 Mbp and had 76 to 94% completion while long read methods recovered genomes between 7.6 Mb to 12.3 Mbp and had completion between 56 to 94% (Fig 4c-d; Fig. S5). The genome recovered by Refmapping for the sample which had the most *Blastocystis* reads (4D, with 2.63 M reads based on Centrifuge results) was found to have the same number of SCGs as the complete ST4 WR1 reference genome (Table S1; Fig 4e). When a second published but smaller and incomplete ST4 genome (BT1) (Table S1) was used as the reference for Refmapping it resulted in genomes that had shorter total length and fewer SCGs across all ST4 samples (Fig 4e-f, Fig S6) further highlighting the dependency of Refmapping on the reference genome used.

### Recovering MRO genomes

We sought to examine whether the reads recovered with Eukfinder and the other methods could also be used to recover mitochondrial genomes from metagenomic datasets. To examine this, we utilized eight real human gut metagenomic datasets to attempt to recover *Blastocystis* mitochondrion-related organelle (MRO) genomes. In 5 samples (3A, 3C, 3D, 4C, and 4D) single complete MRO genomes could be recovered, using identified reads from all methods (Fig 5). In the remaining three samples (3B, 4A, and 4B) the methods recovered no or partial MRO genomes. Among the methods EukRep was unable to recover an MRO genome for sample 3B, and the MRO genomes from the other two samples were smaller than those of the other methods (Fig 5).

**Figure 5.**
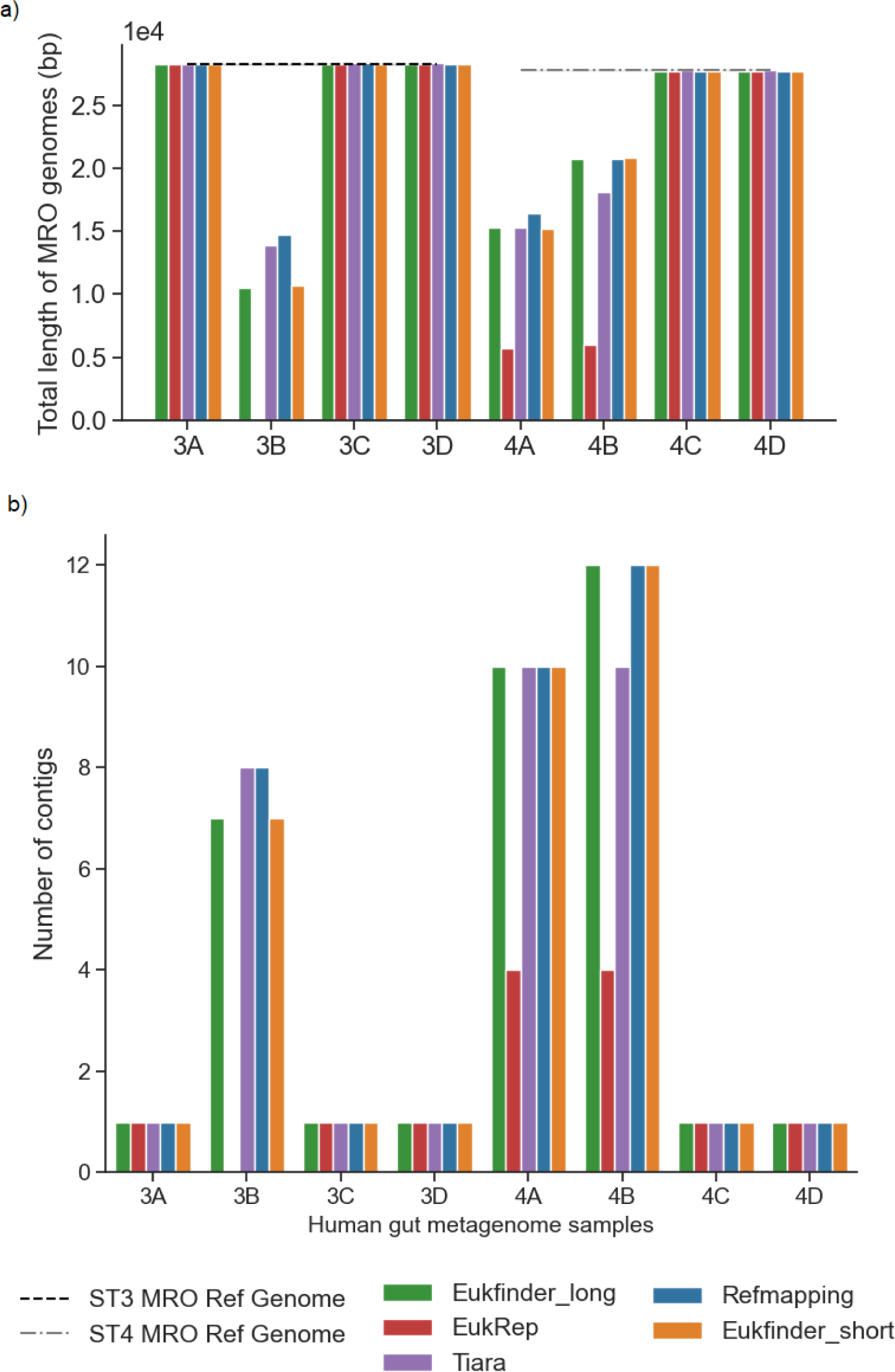
Comparison of Eukfinder to existing methods to recover *Blastocystis* MRO genomes a) Total length of MRO genomes (bp) and b) number of contigs. Eight human gut metagenome datasets were used to compare Eukfinder to EukRep, Refmapping, and Tiara. In three samples (3B, 4A, and 4B), no complete MRO genome was recovered. EukRep completely missed the MRO genome in 3B sample and recovered the second least genomes in the other two samples.

## DISCUSSION

We created the Eukfinder workflow as a solution to recovering nuclear and mitochondrial genomes from eukaryotic data present in metagenomic datasets. Eukfinder improves upon similar bioinformatic tools in several ways. First, Eukfinder is less dependent on reference sequences in comparison to reference mapping methods, allowing for the detection of diverse lineages of microbial eukaryotes. Additionally, Eukfinder can also be used with both short and long sequence reads. Finally, Eukfinder’s databases can be customized to cater to specific environmental samples, further tailoring the program to maximize genome recovery.

Eukfinder_short utilizes short read sequences and provides an initial read classification step that eliminates most prokaryotic sequences prior to assembly. Even though the two Eukfinder methods differed in input data (raw vs assembled reads) and number of rounds of classification (two vs one) Eukfinder_short and Eukfinder_long gave similar results. A previous assessment of the performance of different methods for taxonomic annotation of metagenome data also found that using raw reads or assembled reads yielded similar results (22), however they found that assembly methods were better when the metagenome was large. Looking at the human metagenome samples which ranged from 30-120 M reads we did see slightly larger genome completion with Eukfinder_long and assembled reads when there were >50M reads while below that results were similar.

While Refmapping methods tend to work best for metagenomic samples with a small amount of eukaryotic signal, they are highly dependent on the availability of a high-quality reference genome for the target eukaryote(s), making it not ideal for recovery of genomes from novel species and less-well studied environments. As was seen in this study, even using relatively closely related, but not identical, reference genomes resulted in a greatly reduced recovered genome size of the target eukaryote, as did the use of partial genomes as references. While Eukfinder benefits from the use of high-quality closely-related reference sequences in its databases it can still recover genomes relatively well without them. In this workflow we have implemented a set of parameters for Centrifuge and PLAST, along with specialized databases which maximized our precision and recall when classifying eukaryotic data using Eukfinder. These settings can easily be adjusted however depending on the application or organism of interest. For example, when using metagenomic data to assemble genomes from novel microbial eukaryotes without a reference genome, the minimal hit length, e-value threshold, and coverage cut-off values could be lowered to maximize sequence recovery. In addition, the database can be customized to the environment the sample was taken from. Finally, the supervised binning protocol could also be refined; currently Eukfinder results are combined with information on depth of coverage, Centrifuge identification, PLAST search results against a NCBI nt database, MyCC binning (23) and Metaxa2 (24) results to retrieve eukaryotic contigs.

Aside from MAG discovery from metagenomic data, the Eukfinder workflow can also be applied to tackle other sequencing problems. This pipeline can be used as a decontamination tool for eukaryotic genome sequencing data derived from non-axenic cultures. Such sequencing strategies are common when exploring the genomes of novel protist lineages, for which establishing axenic cultures is cumbersome (25). Applying Eukfinder this way will aid in *de novo* genome assembly of some of the most understudied and poorly understood eukaryotic taxa. Additionally, Eukfinder can be used to pre-screen environmental sequences for depth of coverage of potential eukaryotic community members. Preliminary Eukfinder results can be used in decision making during data collection, such as whether re-sequencing or re-sampling is required. Eukfinder works best if the genome size of interest is <20M and there is a large number of eukaryotic reads, all methods improved with increased number of reads. Given that eukaryotic reads generally make up a small proportion of WGS data in metagenomic samples (typically < 5%), total sequencing reads in the metagenome samples will need to be large to recover complete genomes using Eukfinder. The results of the real human metagenome samples however suggest that it is not just the number of metagenomic or eukaryotic reads that may impact our ability to recover complete genomes. Sample 4D had over 75M metagenomic reads with 3.5% of those being *Blastocystis* (2.6 M reads) and recovered a nearly complete *Blastocystis* ST4 genome, while 3D had nearly 120 M metagenomic reads with just 0.68% of those being *Blastocystis* ST3 (0.82M reads) but was able to recover a genome even more complete than the current reference genome.

In conclusion, these results show that Eukfinder can efficiently generate high-quality near-complete nuclear and mitochondrial genomes from highly diverse microbial eukaryotes such as *Blastocystis*. Eukfinder provides an alternative to the recent tools EukHeist (26) and Whokaryote (27) that build on EukRep and Tiara respectively, with Eukfinder having the added benefit of being able to utilize short raw reads and having increased read recall over both EukRep and Tiara. We anticipate that Eukfinder will be a useful tool for reference-independent and cultivation-free eukaryotic microbial genome recovery from environmental WGS metagenomic sequencing samples.

## MATERIALS AND METHODS

### Implementation of Eukfinder

Eukfinder is a taxonomy-classification-based workflow to recover microbial eukaryotic genomes from WGS metagenomic datasets. A schematic representation of the Eukfinder pipelines is outlined in Fig. 1.

#### (i) Database preparation

The Eukfinder workflow uses two databases: one compatible with Centrifuge (17) and one compatible with PLAST (18). The numbers of genomes in each category are listed in Table S9.

##### Specialized Centrifuge database (DB1)

Centrifuge (17) is a metagenomics taxonomy classification software tool that uses an optimized indexing scheme which allows for the rapid classification of sequencing reads. It contains built-in tools to download genomes from the National Center for Biotechnology Information (NCBI) website and to build custom databases. To maximize Centrifuge’s ability to classify gut metagenome data and eukaryote genomes in particular, a custom database was built (here referred to as specialized DB1). Archaeal, bacterial, and viral genomes associated with the gut microbiome, or without any specific environment listed in the project names, were downloaded from NCBI (as of date). Genomes from all four assembly levels (complete, chromosome, scaffolds, and contigs) were included. An in-house python script was applied to exclude genomes retrieved from environments other than the GI tract (marine, soil, or freshwater). In addition, 4,930 species-level bacterial and archaeal genome bins from >9000 human metagenomes (28) and 913 microbial genomes obtained from rumen metagenomic sequencing (29) were downloaded. Redundant bacterial and archaeal genomes were removed with GTDB-Tk (30) and Treemmer (31). Viral genomes were clustered using MyCC (23) and 40% of the contigs from each cluster (minimum 20 contigs per cluster) were randomly chosen to be included in the database. Eukaryotic genomes from the Kraken2 EupathDB (32) were also included and any pre-downloaded NCBI genomes for those same species were excluded. Additional eukaryotic genomes for protists, fungi, and animals with complete or chromosome level genome assemblies and their corresponding mitochondrial genomes were downloaded from NCBI Genbank (for full list of genomes used see Table S10). In addition, all available genome sequences of *Blastocystis* (Table S1) were also included in DB1, after a decontamination step consisting of mapping the *Blastocystis* genomes against the NCBI nucleotide (nt) database (as of Jan 2019), which did not include any known *Blastocystis* sequences. Contigs in the *Blastocystis* reference genomes that matched >50% of their total length to a bacterium, archaeon, or viral sequence in the nt database and had a nt identity ≥ 80% were considered contaminants and eliminated from the draft genomes used to create specialized DB1. For a full list of excluded *Blastocystis* contigs see Table S11. In-house python scripts were used to build the index files and the centrifuge-build command from Centrifuge was used to construct specialized DB1.

##### Specialized PLAST database (DB2)

PLAST (Parallel Local Alignment Search Tool) (18) is a rapid sequence similarity search tool that is more sensitive though not as fast as Centrifuge. To mitigate computational burden, a specialized PLAST database (hereafter referred to as specialized DB2) was built with a subset of reference genomes from archaea, bacteria, eukaryotic, and mitochondrial genomes selected from the complete set of all the downloaded genomes (Table S12). Specialized DB2 overlaps with specialized DB1 to some degree to enhance the sensitivity of the classification method since PLAST search results were based on similarity (identity) while centrifuge only reports exact alignments with the minimal hit length. For viruses, all viral genomes were downloaded from NCBI Refseq database (ftp.ncbi.nlm.nih.gov/refseq/release/viral/, Mar 2019). All the genome files were combined into a single fasta file, and the database was built using BLAST (33) “makeblastdb” command and a simplified index file containing information that cross-references each sequences accession entry in the database to its respective taxonomic group (i.e., bacteria, archaea, eukaryote, and virus).

### Database with limited or no *Blastocystis*

Comparative genome analysis had revealed that there is a great diversity among the genomes of ST1, ST4 and ST7 and the percentage of unique protein-coding genes ranges from 6% to 20% (Gentekaki et al., 2017), which likely is also the case for other *Blastocystis* genomes in GenBank. Therefore, a combination of all available *Blastocystis* genomes except ST1 (including ST2-ST4, ST6-ST9 genomes, noted as *Blastocystis* ST2-9) was used as reference genomes for the Refmapping method. For Eukfinder, we created two custom Centrifuge and PLAST databases that were either missing the *Blastocystis* ST1 genome (which includes *Blastocystis* ST2-9 genomes) or were missing all *Blastocystis* genomic information. These two databases are referred to as woST1 and woBlasto, respectively.

To examine the performance of Eukfinder_short in recovering genomes when no reference was available, we created two modified versions of both DB1 and DB2. In one modification we created both databases without *Blastocystis* ST1 genome (hereafter referred to as woST1) and in the second modification we removed all *Blastocystis* genomes from DB1 and DB2 (hereafter referred to as woBlasto). We then reran the mock datasets using these databases with Centrifuge and PLAST. In addition, we reran Refmapping replacing the ST1 reference genome with published *Blastocystis* sp. ST2-4, 6-9 as the reference as a parallel test to simulate the situation when there is no direct reference genome.

### Eukfinder pipeline for short sequences

Short reads require a pre-processing step which removes low-quality reads, sequencing adapters, and host reads prior to taxonomic classification with Centrifuge. Eukfinder “read_prep” performs this pre-processing step prior to the first round of classification with Centrifuge. Pre-processing used Trimmomatic v0.36 (34) with a window size of 40 to trim adapters, filter out low-quality bases (<Q25) and remove short reads (<40 bp). In addition, Human host reads were removed by mapping reads to the host reference genome (GRCh38) using Bowtie2 v2.3.1 (38). Processed reads are then assigned taxonomic classification using Centrifuge v1.0.4, using DB1 with a minimum hit length of 40. Two Centrifuge results files (a single file for paired-end reads and a second one for unpaired-end reads) are produced. Eukfinder “short_seqs” uses the pre-processed short read sequence files and Centrifuge results from “read_prep” to separate sequences by taxonomic classification (bacteria, archaea, eukaryote, virus and unknown). Sequences classified as unknown after centrifuge are further analyzed using PLAST (e-value = 0.01, identity = 70% and percent hit length coverage = 30%.), searching against specialized DB2.

After the PLAST analysis all reads classified as either Eukaryotes or Unknown were combined and assembled with metaSPAdes-3.13.1(36) using default parameters (k-mer sizes: auto). All contigs ≥ 1000 bp from the resulting assembly were assigned taxonomic classification using Centrifuge and specialized DB1 and PLAST with specialized DB2. All contigs were given a taxonomic classification (bacteria, archaea, eukaryote, virus and unknown). The contigs labeled Eukaryote (Euk) and Unknown (Unk) (referred collectively as EUnk) were used for supervised binning to recover eukaryotic genome(s).

### Eukfinder pipeline for long sequences

If starting with long-read sequences (Pacbio or Nanopore sequencing reads) or contigs Eukfinder “long_seqs” can be used instead which only requires one round of taxonomic classification using Centrifuge (min hit length =100, DB1) and PLAST (parameters same as “short_seqs”, DB2). Like with Eukfinder_short, contigs are separated into five categories (bacteria, archaea, eukaryote, virus and unknown). Contigs classified as either eukaryote or unknown (EUnk) are used for supervised binning to recover eukaryotic genome(s). All the parameters of Eukfinder used in this paper are listed in Table S13.

### Supervised binning

Supervised binning with MyCC (23) was performed on the contigs classified as either eukaryotic or unknown (i.e., EUnk). Three separate MyCC analyses with different k-mers (4-mer, 5-mer, 5-6-mer) were used to perform binning (Fig. S7). To assist with the final binning, the read-coverage depth for each assembly at each k-mer was calculated by mapping short-reads to contigs with Bowtie2. Following sorting and indexing using SAMtools v1.9 (37) the depth files were generated using MetaBat2 (38) (jgi_summarize_bam_contig_depths). In addition, LSU/SSU rRNA and mitochondrial sequences in the EUnk-assemblies were identified using Metaxa2 (24) with the default database (align length > 300, identity > 90%), and a nucleotide-based PLAST search was conducted using the contigs as queries against the NCBI-NT database (Jan 2019) and the NCBI IDs of the best ‘hits’ were converted to taxonomic IDs with acc2tax v0.6 (github.com/richardmleggett). Read coverage depth, identified rRNA sequences, and taxonomic identities of contigs were mapped to corresponding MyCC bins. For a contig to be included in the final eukaryotic bin (or bins), the following rules were applied:

1. Depth of coverage could not exceed that of the SSU rRNA gene.
2. Best PLAST ‘hit’ could not be a prokaryote or virus with > 90% identity over an aligned length ≥ 1000 bp.
3. A contig was identified as mitochondrial by Metaxa2 and PLAST. (These contigs were marked as mitochondrial genomes.)
4. A contig was identified as eukaryotic by Metaxa2, centrifuge, and /or PLAST.
5. Contigs appeared at least twice in potential eukaryotic clusters across the three MyCC k-mers used.

A schematic of supervised binning is shown in Fig. S8. It is important to mention that, although the supervised binning step is part of the classification workflow, it is not currently implemented in the Eukfinder pipeline, so alternative binning methods (Metabat2 (38) or Anvio (39)) could be used.

### Mock community datasets

To perform a controlled test of Eukfinder capabilities, a series of mock communities were generated by combining all sequence reads from the synthetic ‘Mix-51-staggered’ human gut bacterial data (40; SRR8304765) with randomly selected sets of Illumina sequence reads from the over 34 million reads generated in the published genome of *Blastocystis* sp. ST1 (ATCC 50177/Nand II, GCA_001651215.1). To examine the impact of varying numbers of *Blastocystis* reads on the methods ability to recover the *Blastocystis* genome, eight subsets of reads (ranging from 300K to 10M reads) were randomly selected from among the 34 million Illumina *Blastocystis* reads and combined with all reads from the ‘Mix-51-staggered’ synthetic metagenome dataset. This process was repeated four times for each subset, resulting in 32 mock communities to be analyzed. The total numbers of Illumina ST1 reads, fraction of reads belonging to ST1 and number of contigs ≥ 1000 in each mock community are listed in Table S2.

### Identifying optimal parameters for alternative methods

To test Eukfinder’s capabilities of retrieving *Blastocystis* genomes and examine its performance relative to other eukaryote metagenomic read classifiers we examined genome completeness from each method on both mock community datasets as well as real metagenomic data. To ensure optimal results from all methods, we first identified the parameters for each program that maximized genome completeness and contiguity. We investigated three of the *Blastocystis* read count mock communities (500K, 1M, 2M) to find the best parameters for EukRep, Tiara, and Refmapping methods that would be used for all datasets analyzed.

Refmapping (7) is a method in which metagenomic reads are mapped to the reference genome of an organism of interest. Mapped reads are then assembled, and the resulting contigs are sub-selected for inclusion into the final genome based on length and depth of coverage. Therefore, mapping parameters are important to consider. We tested the Refmapping method using Bowtie2, a memory efficient read mapper, using two different mapping strategies: global (end-to-end) and local mapping. End-to-end mapping relies on a complete alignment of a given read to the reference sequence, while local alignments allow the ends of the reads to be ignored if it increases the overall alignment score.

EukRep (8) classifies metagenomic reads using a machine learning approach that utilizes linear Support Vector Machine (SVM). EukRep has three built in stringency cut-off modes to define how stringent the algorithm is when identifying eukaryotic sequences: strict, balanced, and lenient, from the most to the least stringent. EukRep also has an additional ‘-tie’ flag which specifies the decision for handling sequences when it has an equal number of chunks predicted as eukaryotic and prokaryotic; by default, sequences are classified as eukaryotic. We compared the recovered genomes using EukRep with all three stringency cut-off modes and one analysis under lenient mode with the ‘-tie’ set to prokaryotic.

Tiara (10) utilizes previously trained neural networks to sort metagenomic reads into multiple categories, representing the taxonomic affiliation of each read within a given dataset. Tiara has two main parameters which can be adjusted in its workflow; the k-mer size, which adjusts the size of the DNA substrings used to compute sequence frequency, and the probability threshold, which disregards results with a probability score lower than that. We examined combinations of three different k-mers (4-mer, 5-mer, and 6-mer) with five different probability thresholds (p=0.5, 0.55, 0.6, 0.65, 0.7) on the three mock communities.

### Benchmarking with Mock communities

Eukfinder_short and Refmapping (using the ST1 genome as reference) were used to analyze short reads from all 32 mock communities. Eukfinder_short followed the procedure outlined in Fig. 1a detailed above. For Refmapping, metagenomic short reads were extracted from the dataset by mapping them to the ST1 genome, using Bowtie2 in local mode. All the mapped reads were assembled using metaSPAdes-3.13.1 with default parameters and contigs ≥ 1000 bp were subjected to supervised binning. Resulting bins for both methods were assessed for genome completeness using QUAST. In addition, the short reads were assembled using metaSPAdes-3.13.1 (with default parameters) and all contigs ≥1000 bp were used as input for Eukfinder_long, EukRep, and Tiara for benchmarking. Reads identified as Eukaryotic and/or Unknown by each method were subjected to the same supervised binning procedures and resulting bins were assessed for genome completeness using QUAST.

### Precision and Recall

To assess precision and recall, reads from *Blastocystis* ST1 and ‘Mix-51-staggered’ SRR8304765 were respectively mapped onto contigs for each assembly from the Mock community dataset (‘Mix-51-staggered’ SRR8304765 plus randomly selected *Blastocystis* ST1 Illumina sequencing data) using Bowtie2. Depth of coverage for each contig was calculated using SAMtools v1.9 and depth files were generated using MetaBat2 (jgi_summarize_bam_contig_depths). Contigs having no mapped bacterial reads (zero depth of coverage for SRR8304765 reads) were extracted and classified using centrifuge (database: nt 2019;--min-hitlen 200), Diamond (database: nr Mar2023; −e 0.00001), and Plast (database: nt Mar2022; −e 0.001). All contigs that had ST1 reads mapped and classified to *Blastocystis* were defined as the individual reference genome (IndRef) for each of the 32 mock communities. These IndRef genomes were used as the ground truth for precision and recall analysis. Since Eukfinder_long, EukRep, and Tiara all use the same set of contigs as input, we could assess the precision and recall of these methods to recover those contigs. Precision was measured as the true positives divided by true positives plus false positives. Recall was measured as true positives divided by true positives plus false negatives. True positives were defined as *Blastocystis* contigs correctly identified as *Blastocystis* contigs, and false positives were bacterial contigs identified as *Blastocystis*. True negatives were bacterial contigs identified as bacterial, and false negatives were *Blastocystis* contigs identified as bacterial. The genome fraction recovered by the methods was assessed using QUAST v5.0.2 with two different reference genomes: i) IndRef contigs from each set of randomly subsampled *Blastocystis* reads and ii) the full *Blastocystis* ST1 genome assembly.

### Human gut shotgun metagenomic data

To test Eukfinder’s effectiveness in recovering whole eukaryotic genomes from real metagenomic data, we applied Eukfinder to gut metagenome samples preselected because they were known to contain *Blastocystis*. We identified eight human gut metagenome samples which had > 200,000 reads classified as *Blastocystis* based on Centrifuge results (minimal hit length 30 bp) to test the ability of Eukfinder to reconstruct the *Blastocystis* genome. The eight human gut metagenome samples (Table S7) ranged in total size from 8.8 giga base pairs (Gbp) to 23.3 Gbp and between 48.9 million (M) to 119.7 M raw reads. Between 415K to 2.6M *Blastocystis* reads were identified within these samples using Centrifuge. Eukfinder_short and Refmapping (using either ST3 or ST4 reference genome as appropriate) were used to analyze short reads from the human gut shotgun metagenome samples. In addition, short reads were assembled using metaSPAdes-3.13.1 (with default parameters) and all contigs ≥1000 bp were used as input into Eukfinder_long, EukRep, and Tiara for benchmarking. All eukaryote identified reads from each method underwent the same supervised binning procedures, and resulting bins for all methods were assessed for genome completeness using QUAST and BUSCO.

### Performance measures for eukaryotic genome recovery

We examined Eukfinder (short and long) along with three other methods, Refmapping (7), EukRep (8) and Tiara (10), for their ability to recover the *Blastocystis* genome from the mock communities and human gut metagenome samples. QUAST v5.0.2 (34) was used to assess nuclear genome completeness measured by total genome length and fraction of *Blastocystis* genome recovered. In addition, completeness of the human gut metagenome samples was examined using the number of single copy genes (SCGs) identified using BUSCO 4.0.6 (35) with the stramenophiles_odb10 dataset (21). Performance measures were compared between datasets using pairwise t-tests. In the mock community datasets p-values were corrected for multiple testing using Bonferroni. Performance measures for Eukfinder_short were compared with Refmapping, and performance measures for Eukfinder_long were compared with EukRep, and Tiara.

### Mitochondrial genome recovery

We also assessed whether the Mitochondrial genome could be recovered from within the identified binned reads from each method. We assessed the completeness of the mitochondrial genomes recovered. Eukfinder and EukRep, classify *Blastocystis* mitochondrial reads as Eukaryotic and then during the supervised binning step mitochondrial genomes were identified and separated from nuclear genomes. Refmapping using the mitochondrial reference genomes for ST3 and ST4 (ST3: NC_018042.1, ST4: NC_027962.1) were used to map mitochondrial reads which could then be assembled. Tiara directly classifies reads as mitochondrial, which can then be assembled.

### Statistical analyses

Genome completeness using the different methods on the Mock communities and real human gut metagenomic samples were evaluated using pairwise Student’s t-test (*α* < 0.05). A Bonferroni correction was applied for the mock community datasets.

### Availability of data and materials

Eukfinder is developed and implemented in Python. The source code is posted to the GitHub repository at https://github.com/RogerLab/Eukfinder. Default reference databases can be downloaded from https://perun.biochem.dal.ca/Metagenomics-Scavenger/

## SUPPLEMENTAL MATERIAL

Supplemental material for this article is available online only.

## ACKNOWLEDGEMENTS

This work was supported by a Canadian Institutes for Health Research Foundation Grant FRN-142349 awarded to A.J.R. Computations were performed in part on the Niagara supercomputer at the SciNet HPC Consortium. SciNet is funded by Innovation, Science and Economic Development Canada; the Digital Research Alliance of Canada; the Ontario Research Fund: Research Excellence; and the University of Toronto.

We declare that we have no competing interests.

---

**Fig S1.**
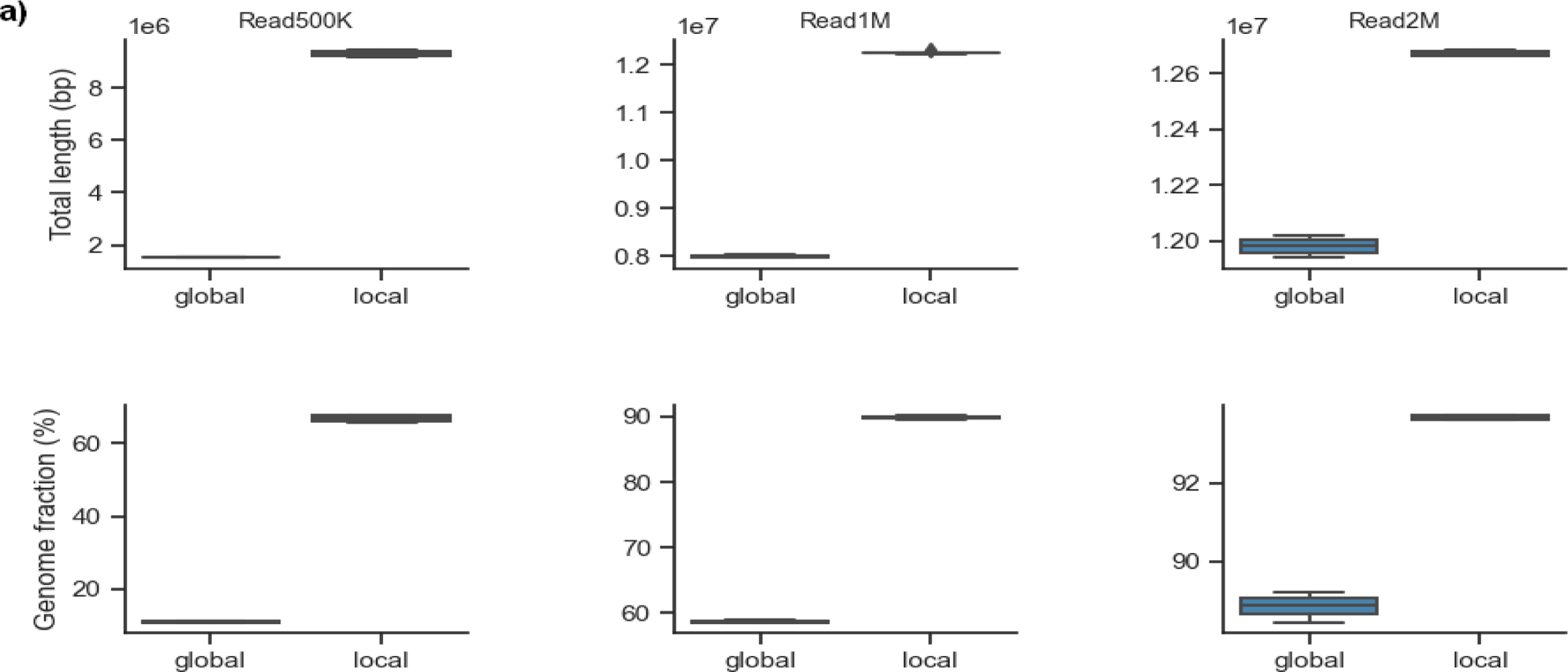
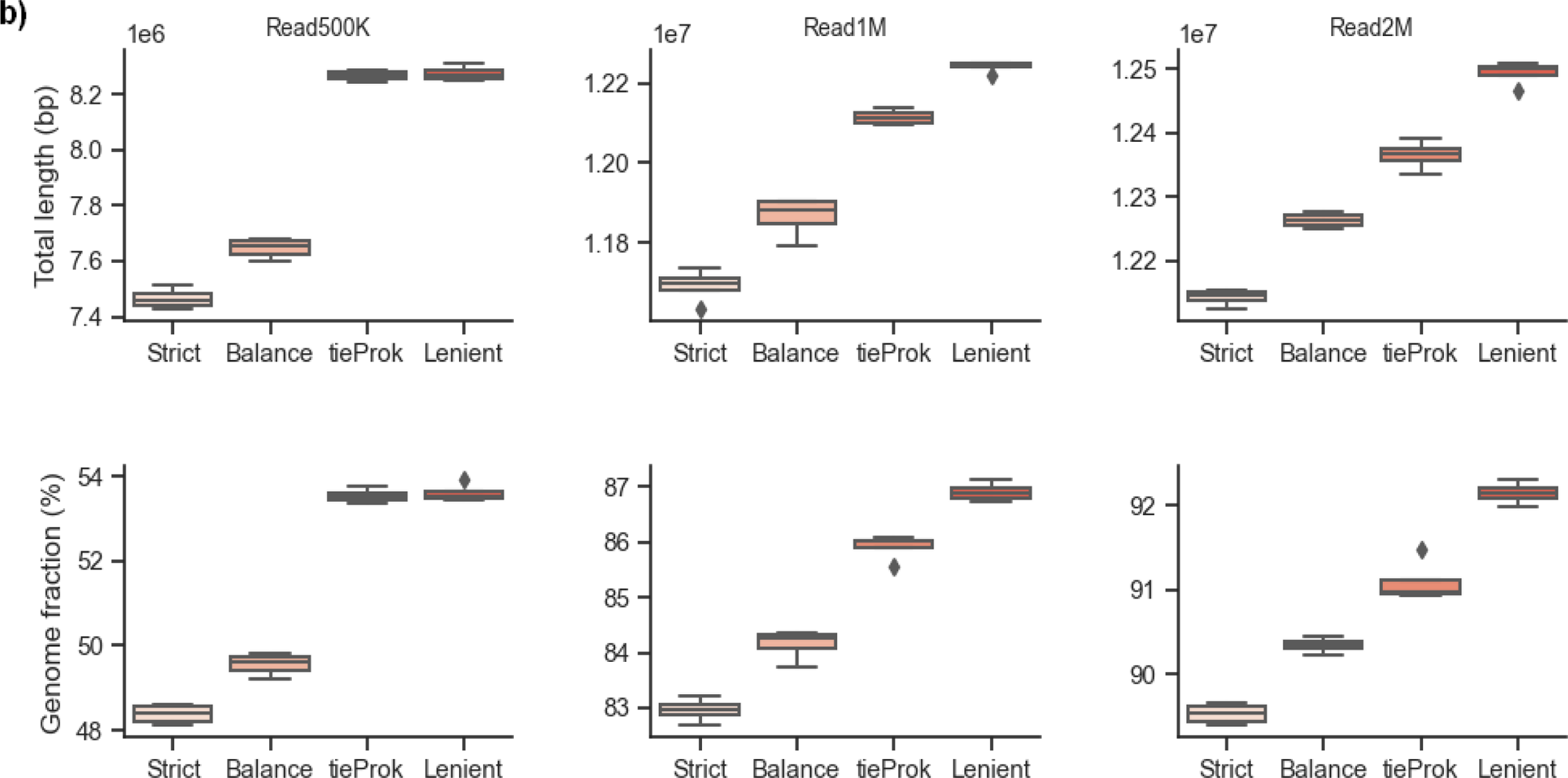
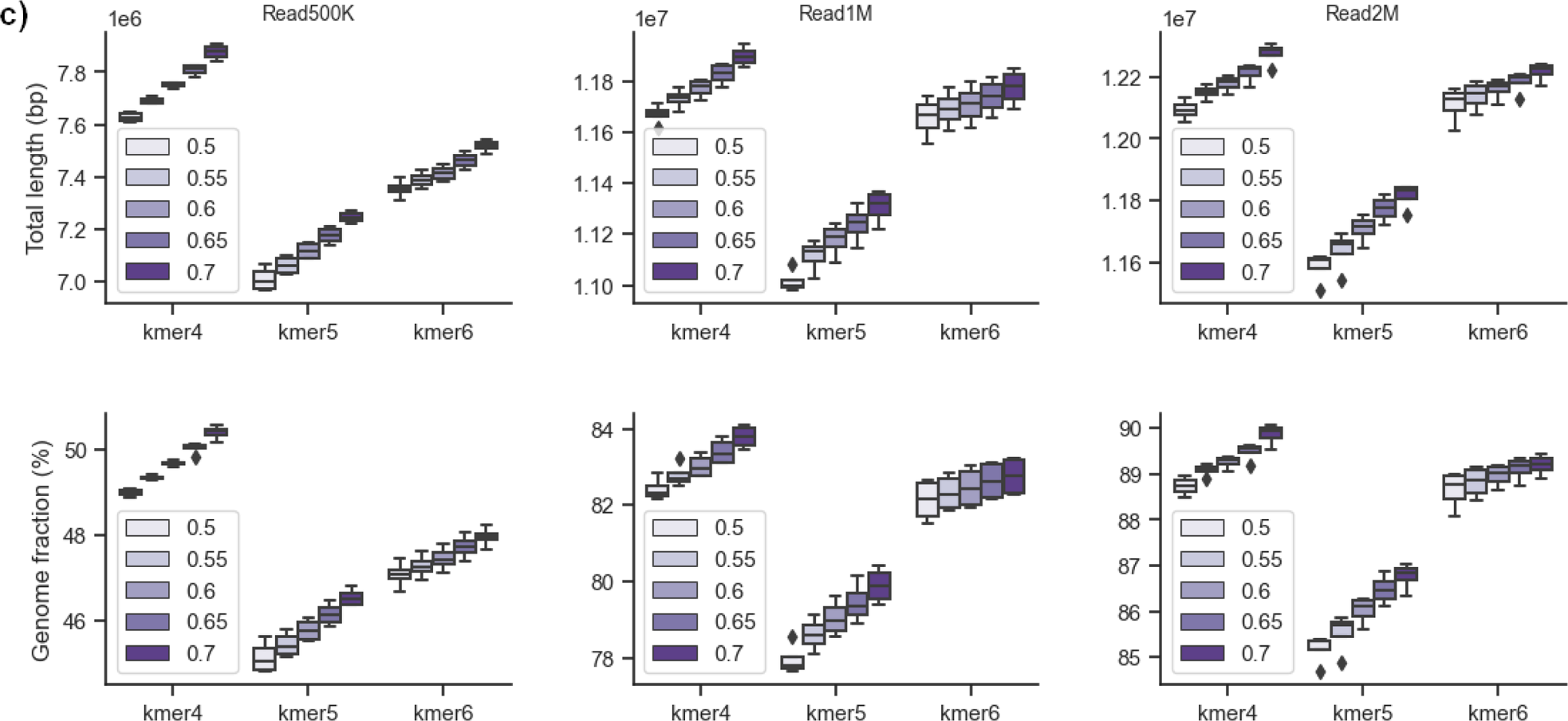
Plots of the effects of different parameters for Refmapping, EukRep, and Tiara, in the recovery of Blastocystis ST1 total length (bp) and genome fraction (%) over three different Blastocystis read count mock community datasets (500K, 1M, and 2M) to identify optimal run parameters. a) The effects of local and global mode for reference mapping were examined in Refmapping. b) Three algorithm stringency cut-off modes were examined (strict, balanced, and lenient) as well as the “tie” parameter, which resolves how ties are placed, eukaryote (default) or prokaryote in the program EukRep. c) Three k-mer sizes (4-mer, 5-mer, 6-mer) and five different probability thresholds (p=0.5, 0.55, 0.6, 0.65, 0.7) were examined in Tiara.

**Fig. S2.**
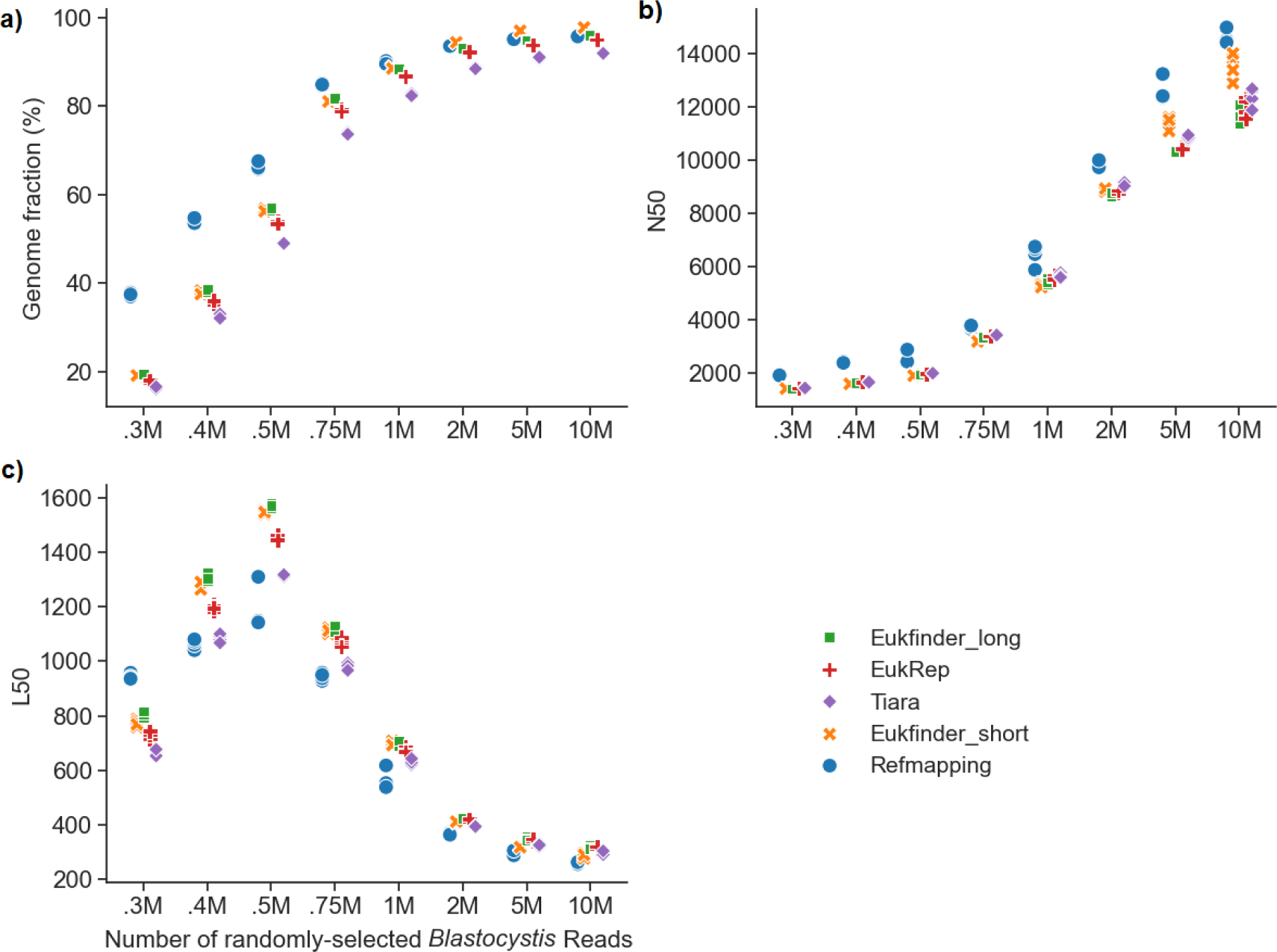
Comparison of methods (Eukfinder_short, Refmapping, Eukfinder_long, EukRep, and Tiara) to recover *Blastocystis* ST1 genomes in eight datasets of bacterial mock metagenomic data (staggered mix-51) and varying amounts of randomly selected *Blastocystis* ST1 Illumina sequencing reads. Plot of *Blastocystis* ST1 QUAST assessed a) genome fraction (%), b) N50 and c) L50 for each of the 8 mock community datasets. EukRep was run in lenient mode, Tiara was run with kmer of 4 and probability threshold of 0.7. Refmapping was run in local mode using *Blastocystis* ST1 genome as the reference genome.

**Fig. S3.**
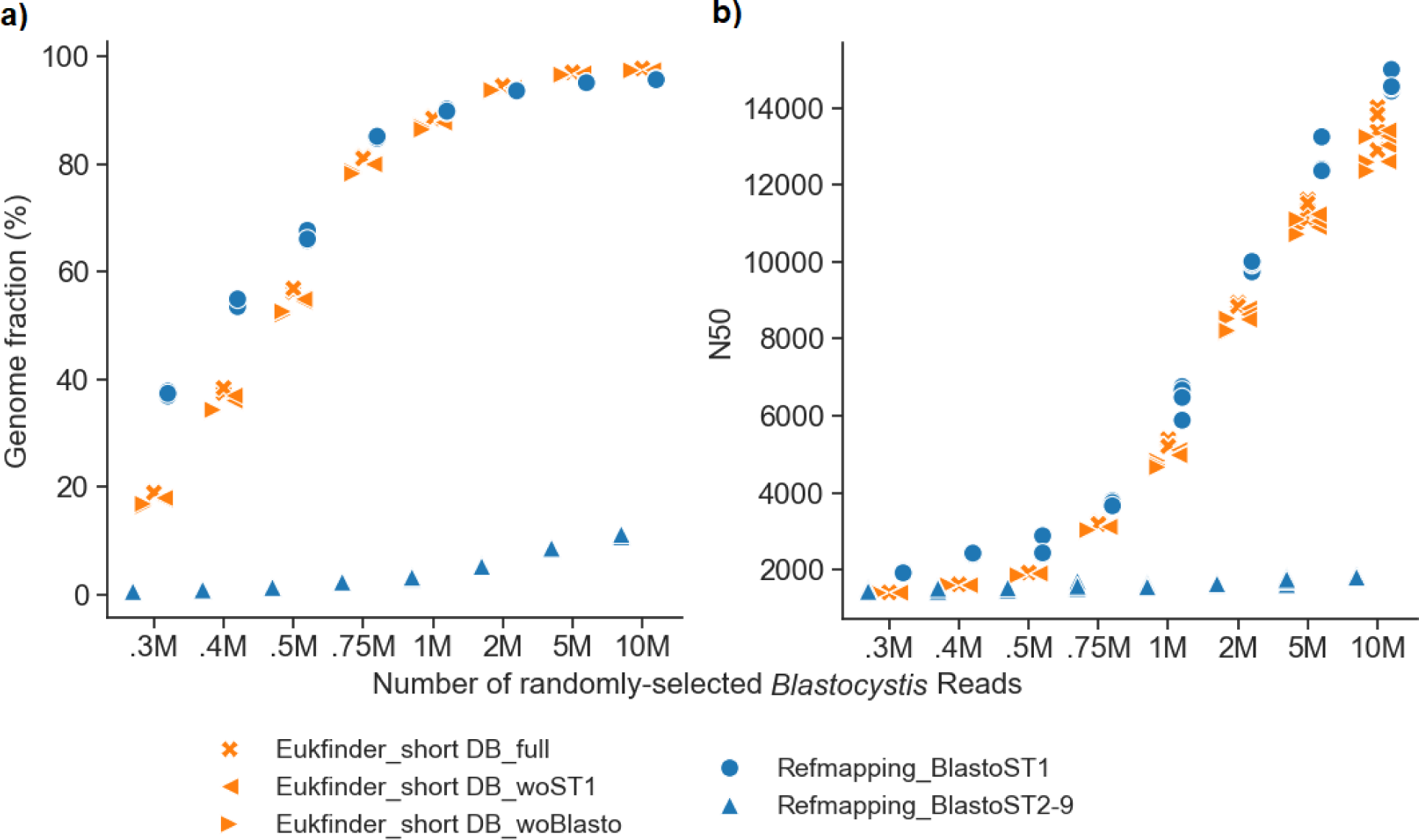
Comparison of Eukfinder_short to Refmapping with or without the aid of the published ST1 genome, based on a) genome fraction recovered (%) and b) N50 by QUAST when compared to the given reference genome. For Eukfinder, three sets of databases were used: the complete database (noted as full), the database missing the *Blastocystis* ST1 genome (which includes *Blastocystis* ST2-ST4, ST6-ST9 genomes, noted as woST1) and the one missing all *Blastocystis* genomic information (noted as woBlasto). For the Refmapping method, ST1 or a combination of all available *Blastocystis* genomes except ST1 (noted as BlastoST2-9) was used as reference genome.

**Fig S4.**
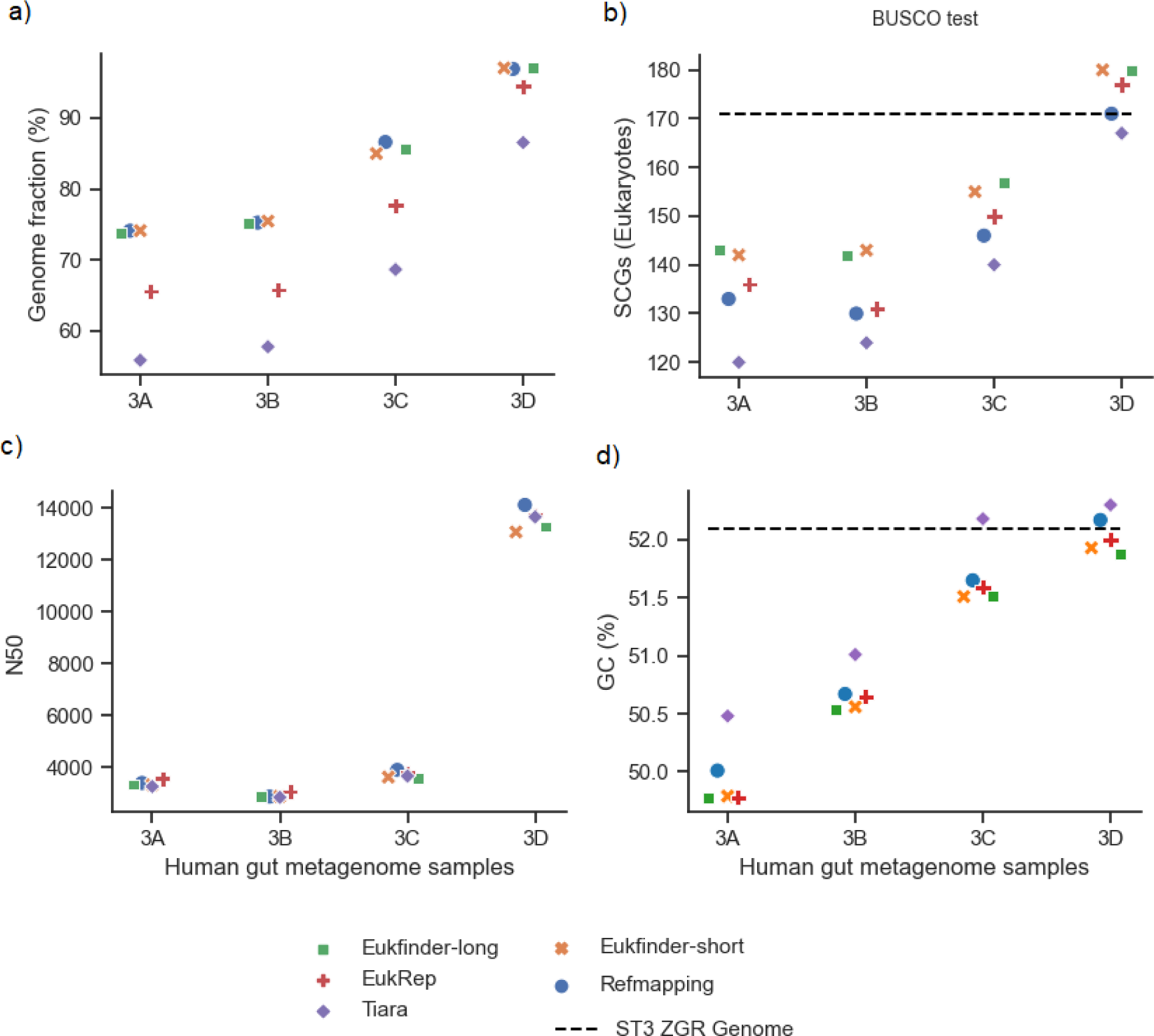
*Blastocystis* ST3 genomes recovered from four human gut metagenome samples (3A-D) using Refmapping, Eukfinder_short, Eukfinder_long, EukRep and Tiara. Completeness of ST3 genomes in samples based on a) genome fraction recovered (%) assessed using QUAST; b) number of single-copy genes (SCGs) detected using BUSCO (eukaryote_odb10); c) N50 assessed using QUAST and d) percent GC content. Tiara was run with kmer of 4 and probability threshold of 0.7. Refmapping was run in local mode using *Blastocystis* ST3 ZGR as the reference genome.

**Fig S5.**
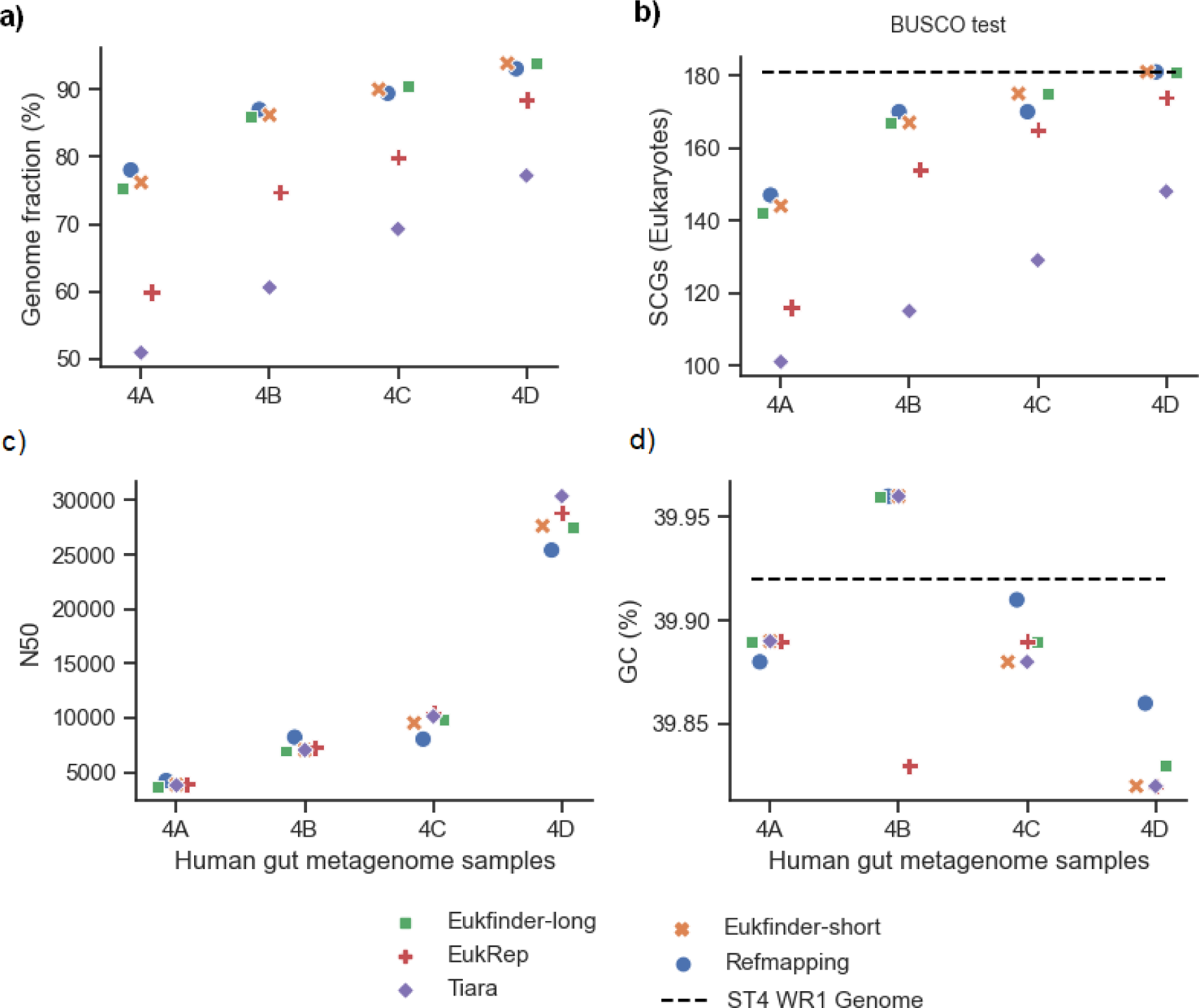
*Blastocystis* ST4 genomes recovered from four human gut metagenome samples (4A-D) using Refmapping, Eukfinder_short, Eukfinder_long, EukRep and Tiara. Completeness of ST4 genomes in samples based on a) genome fraction recovered (%) assessed using QUAST; b) number of single-copy genes (SCGs) detected using BUSCO (eukaryote_odb10); c) N50 and d) percent GC content assessed using QUAST. EukRep was using lenient, Tiara was run with kmer of 4 and probability threshold of 0.7 and Refmapping was run in local mode using *Blastocystis* ST4 WR1 (black dashed line) and BT1 gray dash-dot line as the reference genomes.

**Fig S6.**
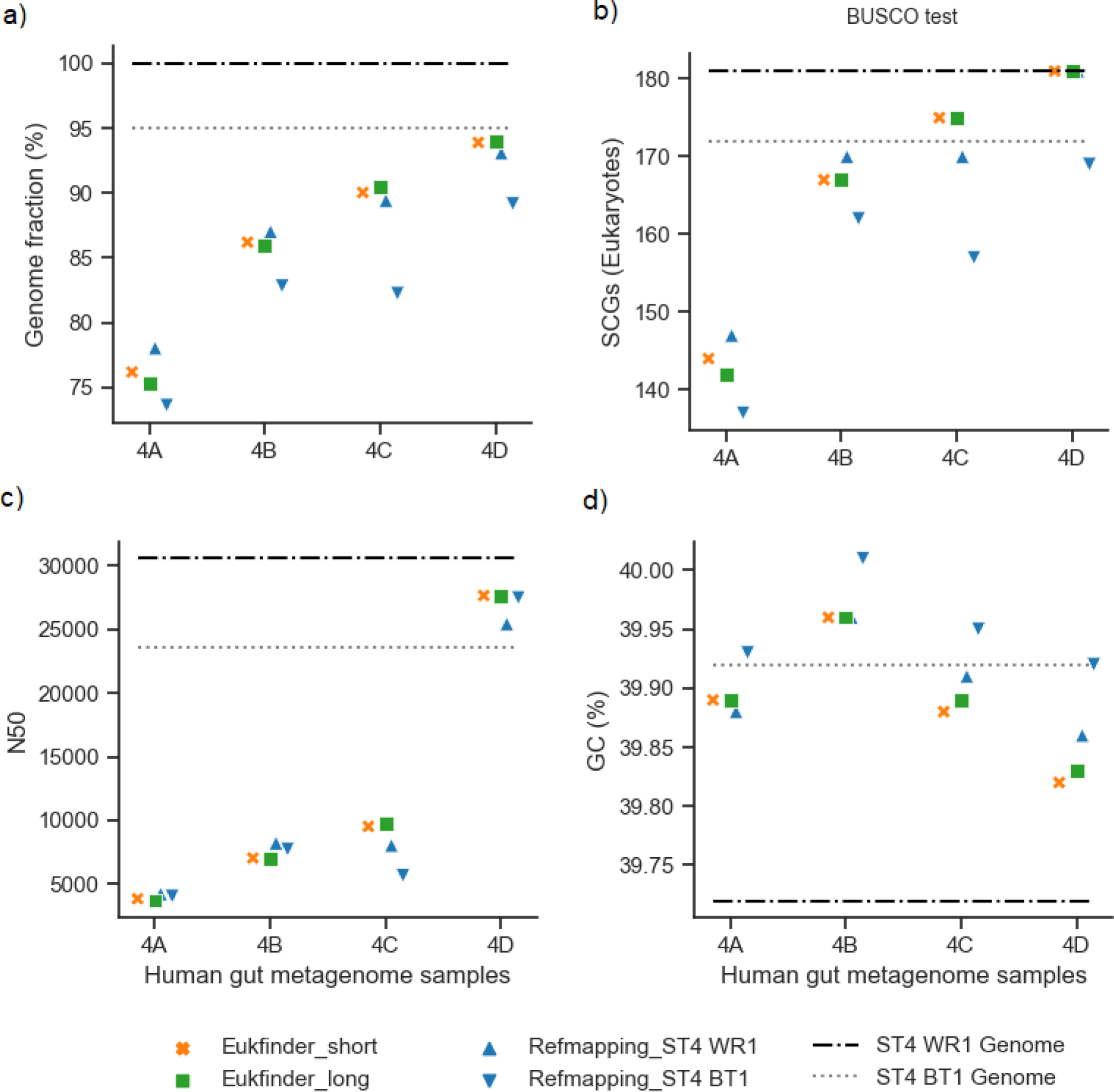
*Blastocystis* ST4 genomes recovered from four human gut metagenome samples (4A-D) using Refmapping, Eukfinder_short, Eukfinder_long, EukRep and Tiara. Completeness of ST4 genomes in samples based on a) genome fraction recovered (%) assessed using QUAST; b) number of single-copy genes (SCGs) detected using BUSCO (eukaryote_odb10); c) N50 and d) percent GC content assessed using QUAST. EukRep was using lenient, Tiara was run with kmer of 4 and probability threshold of 0.7 and Refmapping was run in local mode using *Blastocystis* ST4 WR1 or BT1 as the reference genomes.

**Fig. S7.**
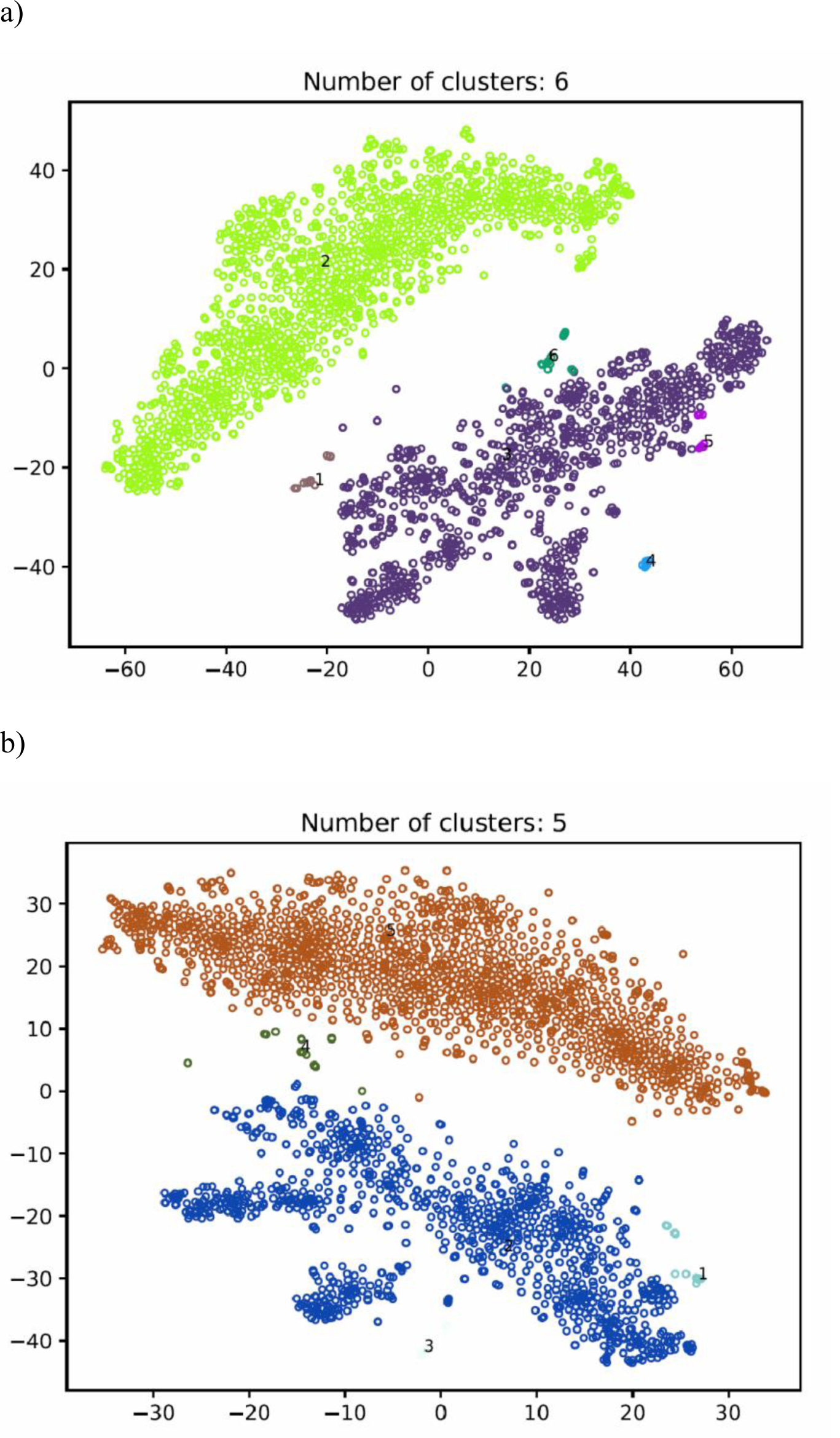
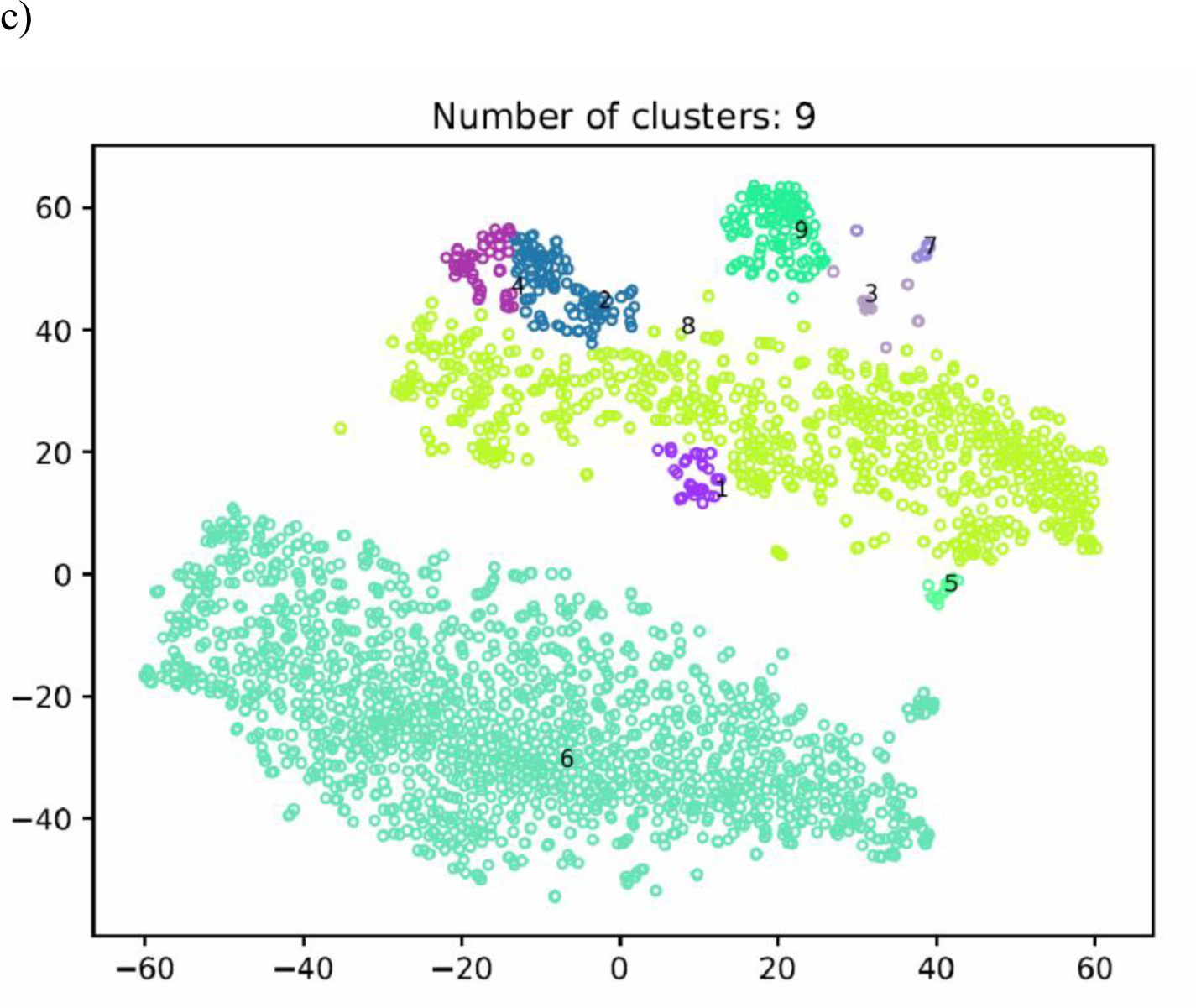
Example of Genome cluster maps generated by MyCC using three k-mers: a) 4mer, b) 5mer, and c) 56mer. Each circle represents a contig. Each cluster is shown in a different color and represents a possible genome.

**Fig. S8.**
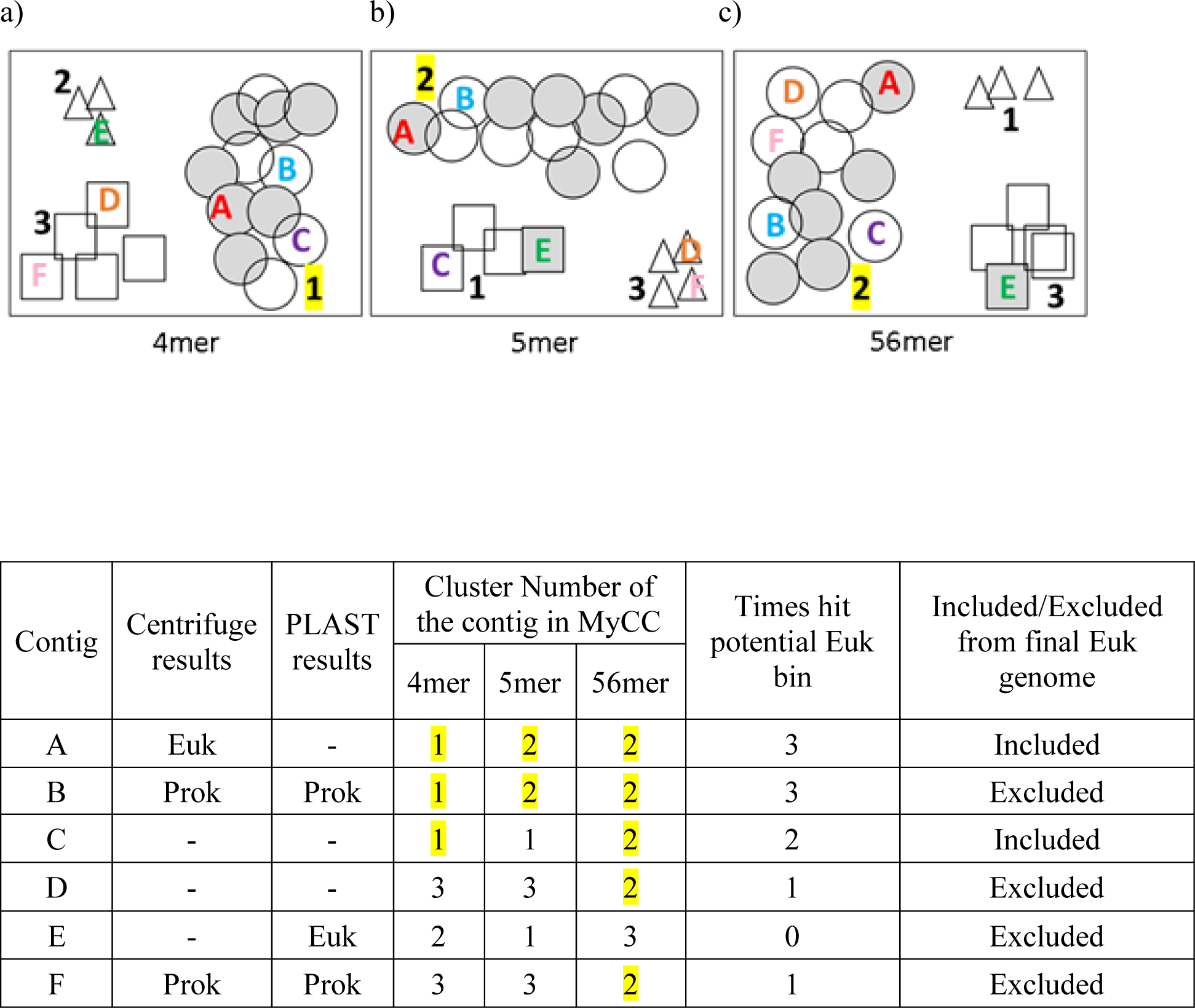
The schematic explanation of how eukaryotic contigs are selected based on MyCC binning, Centrifuge, and PLAST results. a) –c) represent the plots of cluster maps generated by MyCC based on marker genes, k-mer usage and depth of coverage for each k-mer (marked under the box). The geometric shapes (triangles, squares, and circles) represent contigs in different clusters. Contigs with a hit to eukaryotes by Centrifuge or PLAST are shaded in gray. Digital numbers in each plot represent the cluster number. The numbers of potential eukaryotic clusters are highlighted yellow. The alphabet letters A – E represent the contigs that appeared at least once in the potential eukaryotic clusters. To be included in a eukaryotic genome, a contig must appear in at least twice in the potential eukaryotic clusters across different values of k-mers (Contigs A-C). Note that 56mer represents a combination of 5mer and 6mer.

**Table S1.** Genomic features of published *Blastocystis* reference genomes. Subtypes in bold are complete or near complete genomes while all others are incomplete.

| *Blastocystis*<br>subtype &<br>isolate | GenBank<br>Accession<br>Number | Size<br>(Mb) | Scaffolds | GC<br>content<br>(%) | Assembly<br>Level | Year<br>published | Reference |
| --- | --- | --- | --- | --- | --- | --- | --- |
| **ST1**<br>**Nand II** | **GCA_001651215** | 16.4683 | 580 | 53.00 | Scaffold | 2017 | Gentekaki et al., 2017 |
| ST2<br>Flemming | GCA_000963365 | 12.6931 | 969 | 54.00 | Scaffold | 2015 | Andersen et al., 2015 |
| ST3 ZGR | GCA_000963385 | 11.6514 | 917 | 52.00 | Scaffold | 2015 | Andersen et al., 2015 |
| **ST4 WR1** | **GCA_000743755** | 12.9194 | 1301 | 39.70 | Scaffold | 2015 | Wawrzyniak et al., 2015 |
| ST4 BT1 | GCA_000963395 | 11.5409 | 849 | 39.90 | Scaffold | 2015 | Andersen et al., 2015 |
| ST6<br>SSI:754 | GCA_000963415 | 15.4178 | 879 | 43.10 | Scaffold | 2015 | Andersen et al., 2015 |
| **ST7**<br>**isolate B** | **GCA_000151665** | 18.8172 | 54 | 45.30 | Scaffold | 2011 | Denoeud et al., 2011 |
| ST7 ASY-1 | GCA_003575125 | 10.4299 | 10257 | 52.00 | Contig | 2018 | NCBI<br>PRJNA482946 |
| ST8 Dmp/<br>08-128 | GCA_000963455 | 12.2390 | 947 | 39.70 | Scaffold | 2015 | Andersen et al., 2015 |
| ST9 F5323 | GCA_000963465 | 11.7149 | 871 | 43.00 | Scaffold | 2015 | Andersen et al., 2015 |

**Table S2.** Numbers of randomly selected *Blastocystis* ST1 Illumina sequencing reads that were combined with the synthetic ‘Mix-51-staggered’ human gut bacterial data of Miller et al (2019, SRR8304765) to create the mock metagenomic datasets used in this study.

| **Identifier**<br>**(# Reads)** | **Number of**<br>***Blastocystis* reads** | **Total Number of**<br>**Reads (M)** | **Fraction of total**<br>**dataset (%)** |
| --- | --- | --- | --- |
| 300K | 300,000 | 16.9 | 1.78 |
| 400K | 400, 000 | 17 | 2.35 |
| 500K | 500, 000 | 17.1 | 2.92 |
| 750K | 750, 000 | 17.35 | 4.32 |
| 1M | 1, 000, 000 | 17.6 | 5.68 |
| 2M | 2, 000, 000 | 18.6 | 10.75 |
| 5M | 5, 000, 000 | 21.6 | 23.15 |
| 10M | 10, 000, 000 | 26.6 | 37.59 |

**Table S4.** Pairwise Student’s t-test comparisons of total recovered length and genome fraction of the simulated mock metagenome communities (# Reads: numbers of randomly selected *Blastocystis* ST1 Illumina sequencing reads, 300K to 10M) using Eukfinder_short and Refmapping (with *Blastocystis* ST1 reference genome). Bonferroni correction for multiple tests was applied. The null hypothesis tested was pairwise mean total length or genome fraction of Eukfinder_short was ≤ Refmapping and the alternative was that the pairwise mean total length or genome fraction of Eukfinder_short was > Refmapping.

| # Reads | Total Length (bp) |  |  | Genome Fraction(%) |  |  |
| --- | --- | --- | --- | --- | --- | --- |
|  | t_stat | p_value | Bonferroni corrected p-value | t_stat | p_value | Bonferroni corrected p-value |
| 300K | -97.4 | 1 | 1 | -119.5 | 1 | 1 |
| 400K | -87.2 | 1 | 1 | -123 | 1 | 1 |
| 500K | -13.4 | 1 | 1 | -22.4 | 1 | 1 |
| 750K | -9.4 | 1 | 1 | -19 | 1 | 1 |
| 1M | -4.6 | 0.99 | 1 | -11.0 | 1 | 1 |
| 2M | 12.2 | **5.96e-4** | **0.005** | 19.4 | **1.50e-4** | **0.0012** |
| 5M | 17.6 | **1.99e-4** | **0.002** | 46.8 | **1.08e-5** | **8.64e-5** |
| 10M | 40.8 | **1.62e-5** | **1.30e-4** | 36.7 | **2.22e-5** | **1.78e-4** |

**Table S5.** Pairwise Student’s t-test comparisons of total recovered length and genome fraction of the simulated mock metagenome communities using Eukfinder_longer and Eukrep (# Reads: numbers of randomly selected *Blastocystis* ST1 Illumina sequencing reads, 300K to 10M). Bonferroni correction for multiple tests was applied. The null hypothesis tested was pairwise mean total length or genome fraction of Eukfinder_long was ≤ Eukrep and the alternative was that the pairwise mean total length or genome fraction of Eukfinder_long was > Eukrep.

| #<br>Reads | Total Length (bp) |  |  | Genome Fraction (%) |  |  |
| --- | --- | --- | --- | --- | --- | --- |
|  | t_stat | p_value | Bonferroni corrected<br>p-value | t_stat | p_value | Bonferroni corrected<br>p-value |
| 300K | 40 | **1.72e-05** | **1.38e-04** | 41.8 | **1.51e-05** | **1.21e-04** |
| 400K | 46.5 | **1.09e-05** | **8.76e-05** | 56.3 | **6.17e-06** | **4.94e-05** |
| 500K | 34.4 | **2.70e-05** | **2.16e-04** | 33.5 | **2.93e-05** | **2.34e-04** |
| 750K | 19.4 | **1.58e-04** | **1.27e-03** | 18.1 | **1.85e-04** | **0.001** |
| 1M | 59.5 | **5.24e-06** | **4.19e-05** | 53.7 | **7.13e-06** | **5.70e-05** |
| 2M | 18.7 | **1.67e-04** | **1.34e-03** | 20.3 | **1.30e-04** | **0.001** |
| 5M | 20.7 | **1.23e-04** | **9.83e-04** | 20.2 | **1.32e-04** | **0.001** |
| 10M | 18.3 | **1.77e-04** | **1.41e-03** | 17.1 | **2.17e-04** | **0.002** |

**Table S6.** Pairwise Student’s t-test comparisons of total recovered length and genome fraction of the simulated mock metagenome communities using Eukfinder_longer and Tiara (# Reads: numbers of randomly selected *Blastocystis* ST1 Illumina sequencing reads, 300K to 10M). Bonferroni correction for multiple tests was applied. The null hypothesis tested was pairwise mean total length or genome fraction of Eukfinder_long was ≤ Tiara and the alternative was that the pairwise mean total length or genome fraction of Eukfinder_long was > Tiara.

| #<br>Reads | Total Length (bp) |  |  | Genome Fraction(%) |  |  |
| --- | --- | --- | --- | --- | --- | --- |
|  | t_stat | p_value | Bonferroni corrected<br>p-value | t_stat | p_value | Bonferroni corrected<br>p-value |
| 300K | 41.8 | **1.50E-05** | **1.20E-04** | 38.5 | **1.92E-05** | **1.54E-04** |
| 400K | 128.1 | **5.24E-07** | **4.19E-06** | 142.4 | **3.82E-07** | **3.05E-06** |
| 500K | 113.8 | **7.48E-07** | **5.98E-06** | 93.8 | **1.33E-06** | **1.07E-05** |
| 750K | 113.8 | **7.48E-07** | **5.98E-06** | 106 | **9.25E-07** | **7.40E-06** |
| 1M | 54.4 | **6.84E-06** | **5.47E-05** | 57 | **5.95E-06** | **4.76E-05** |
| 2M | 35.9 | **2.37E-05** | **1.90E-04** | 35.3 | **2.50E-05** | **2.00E-04** |
| 5M | 32.8 | **3.11E-05** | **2.49E-04** | 33.5 | **2.93E-05** | **2.34E-04** |
| 10M | 68.9 | **3.37E-06** | **2.70E-05** | 60.7 | **4.92E-06** | **3.94E-05** |

**Table S7.** General description of the tested human gut metagenome datasets.

| Name | Dataset | Size (Gbp) | Number of Total Reads (M) | Assembly (Mb) | *Blastocystis* subtype | *Blastocystis* Reads (M) | Percentage of total dataset | Reference |
| --- | --- | --- | --- | --- | --- | --- | --- | --- |
| 3A | SRR061218 | 6.0 | 30.2 | 141.0 | ST3 | 0.50 | 1.67% | Lloyd-Price J et al. 2017 |
| 3B | SRR060363 | 7.7 | 37.9 | 113.4 | ST3 | 0.48 | 1.27% | Lloyd-Price J et al. 2017 |
| 3C | ERR636351 | 11.1 | 56.7 | 285.7 | ST3 | 0.42 | 0.74% | Bengtsson-Palme J et al. 2015 |
| 3D | ERR636414 | 23.3 | 119.7 | 319.4 | ST3 | 0.82 | 0.68% | Bengtsson-Palme J et al. 2015 |
| 4A | ERR321632 | 8.5 | 47.2 | 240.9 | ST4 | 0.45 | 0.95% | Le Chatelier E et al. 2013 |
| 4B | ERR321638 | 8.7 | 48.4 | 201.0 | ST4 | 0.53 | 1.09% | Le Chatelier E et al. 2013 |
| 4C | ERR636359 | 16.5 | 84.2 | 398.7 | ST4 | 0.59 | 0.70% | Bengtsson-Palme J et al. 2015 |
| 4D | ERR636397 | 13.6 | 75.3 | 294.1 | ST4 | 2.63 | 3.4% | Bengtsson-Palme J et al. 2015 |

**Table S9.**
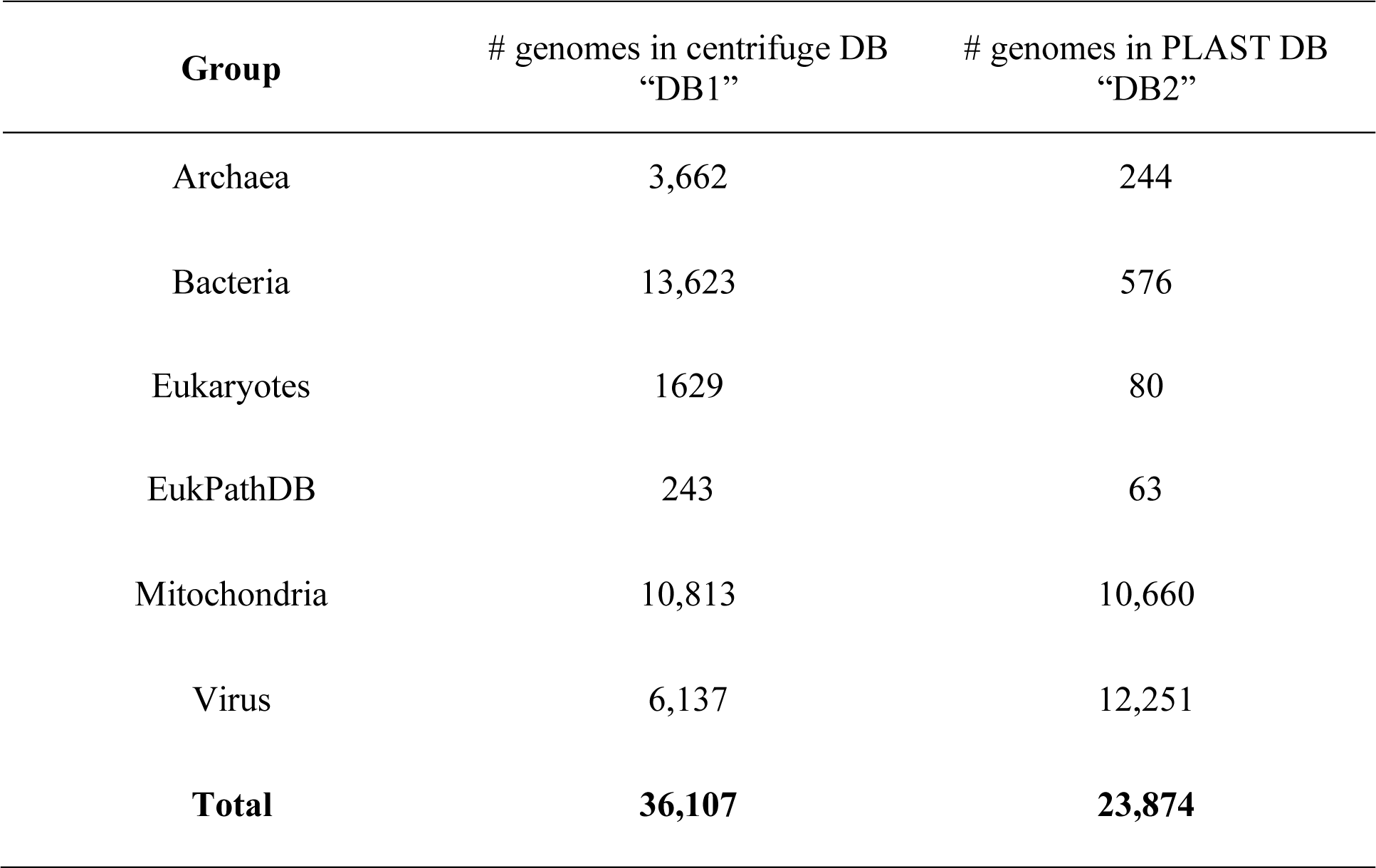
Numbers of genomes present in the specialized databases by taxonomic group.

| **Group** | **# genomes in centrifuge DB**<br>**“DB1”** | **# genomes in PLAST DB**<br>**“DB2”** |
| --- | --- | --- |
| Archaea | 3,662 | 244 |
| Bacteria | 13,623 | 576 |
| Eukaryotes | 1629 | 80 |
| EukPathDB | 243 | 63 |
| Mitochondria | 10,813 | 10,660 |
| Virus | 6,137 | 12,251 |
| **Total** | **36,107** | **23,874** |

**Table S11.** Contigs removed from *Blastocystis* reference genomes. MRO genomes are labelled with asterisks.

| *Blastocystis* Subtype | Contig | *Blastocystis* Subtype | Contig |
| --- | --- | --- | --- |
| ST2 | JZRJ01000088 * | ST6 | JZRM01000240 |
| ST2 | JZRJ01000923 | ST6 | JZRM01000270 |
| ST3 | JZRK01000047 * | ST6 | JZRM01000285 |
| ST3 | JZRK010000726 | ST6 | JZRM01000317 |
| ST3 | JZRK01000382 | ST6 | JZRM01000353 |
| ST3 | JZRK01000623 | ST6 | JZRM01000399 |
| ST3 | JZRK01000726 | ST6 | JZRM01000403 |
| ST3 | JZRK01000754 | ST6 | JZRM01000411 |
| ST3 | JZRK01000820 | ST6 | JZRM01000419 |
| ST3 | JZRK01000826 | ST6 | JZRM01000431 |
| ST3 | JZRK01000839 | ST6 | JZRM01000456 |
| ST6 | JZRM01000006 | ST6 | JZRM01000464 |
| ST6 | JZRM01000011 | ST6 | JZRM01000480 |
| ST6 | JZRM01000013 | ST6 | JZRM01000507 |
| ST6 | JZRM01000016 | ST6 | JZRM01000535 |
| ST6 | JZRM01000019 | ST6 | JZRM01000542 |
| ST6 | JZRM01000022 | ST6 | JZRM01000570 |
| ST6 | JZRM01000023 | ST6 | JZRM01000598 |
| ST6 | JZRM01000027 | ST6 | JZRM01000617 |
| ST6 | JZRM01000029 | ST6 | JZRM01000659 |
| ST6 | JZRM01000030 | ST6 | JZRM01000755 |
| ST6 | JZRM01000035 * | ST6 | JZRM01000790 |
| ST6 | JZRM01000036 | ST6 | JZRM01000826 |
| ST6 | JZRM01000052 | ST6 | JZRM01000830 |
| ST6 | JZRM01000058 | ST6 | JZRM01000879 |
| ST6 | JZRM01000081 | ST8 | JZRN01000022 * |
| ST6 | JZRM01000084 | ST8 | JZRN01000233 |
| ST6 | JZRM01000095 | ST8 | JZRN01000747 |
| ST6 | JZRM01000101 | ST8 | JZRN01000879 |
| ST6 | JZRM01000108 | ST9 | JZRO01000015 * |
| ST6 | JZRM01000117 | ST9 | JZRO01000142 |
| ST6 | JZRM01000122 | ST9 | JZRO01000228 |
| ST6 | JZRM01000123 | ST9 | JZRO01000234 |
| ST6 | JZRM01000137 | ST9 | JZRO01000235 |
| ST6 | JZRM01000150 | ST9 | JZRO01000417 |
| ST6 | JZRM01000163 | ST9 | JZRO01000444 |
| ST6 | JZRM01000189 | ST9 | JZRO01000788 |
| ST6 | JZRM01000198 | ST9 | JZRO01000789 |
| ST6 | JZRM01000214 | ST9 | JZRO01000859 |
| ST6 | JZRM01000230 | ST9 | JZRO01000871 |

**Table S13.**
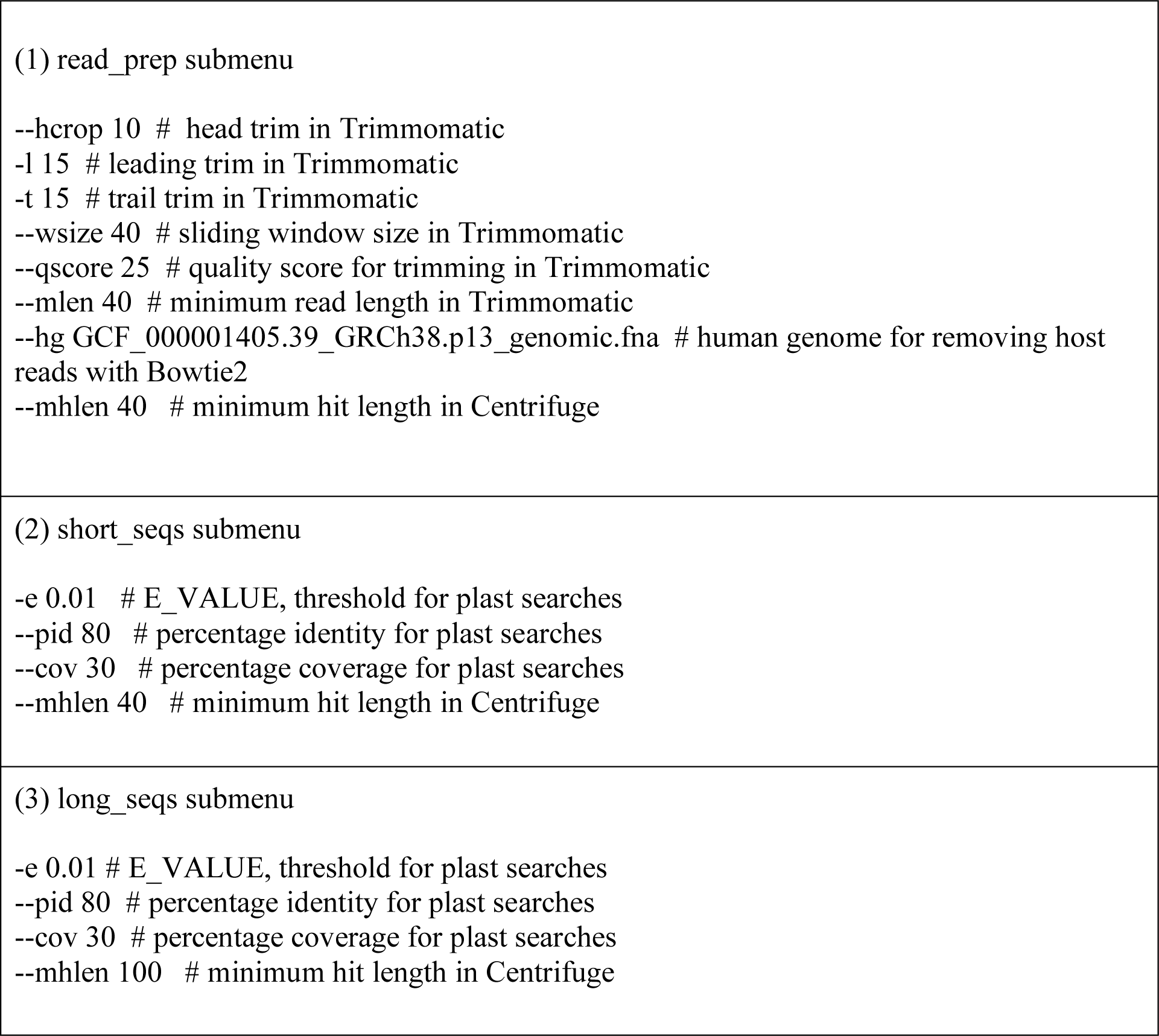
Parameters of Eukfinder used in this paper. Specific information corresponding for each flag is provided after the hashtag.

